# Kindlin-2 preserves integrity of the articular cartilage to protect against osteoarthritis

**DOI:** 10.1101/2021.08.11.456023

**Authors:** Xiaohao Wu, Yumei Lai, Sheng Chen, Chunlei Zhou, Chu Tao, Xuekun Fu, Jun Li, Jian Huang, Wei Tong, Hongtao Tian, Zengwu Shao, Chuanju Liu, Di Chen, Xiaochun Bai, Huiling Cao, Guozhi Xiao

## Abstract

Osteoarthritis (OA) is an aging-related degenerative joint disease, which has no cure partly due to limited understanding of its pathological mechanism(s). Here we report that the focal adhesion protein Kindlin-2, but not Kindlin-1 or −3, is highly expressed in articular chondrocytes of the hyaline cartilage, which is dramatically decreased in the degenerated articular cartilage of aged mice and patients with OA. Inducible deletion of Kindlin-2 in chondrocytes at adult stage leads to spontaneous OA and much severe OA lesions in the mice receiving the surgery of destabilization of the medial meniscus. Mechanistically, Kindlin-2 deficiency promotes mitochondrial oxidative stress and activates Stat3 in articular chondrocytes, leading to Runx2-mediated chondrocyte hypertrophic differentiation and catabolism. In vivo, systemic pharmacological blockade of Stat3 activation or genetic ablation of Stat3 in chondrocytes reverses aberrant accumulation of Runx2 and ECM-degrading enzymes and limits OA deteriorations caused by Kindlin-2 deficiency. Furthermore, genetic inactivation of Runx2 in chondrocytes reverses structural changes and OA lesions caused by Kindlin-2 deletion without down-regulating p-Stat3 in articular chondrocytes. Of translational significance, intraarticular injection of Kindlin-2-expressing adeno-associated virus decelerates progression of aging- and instability-induced knee joint OA in mice. Collectively, we identify a novel pathway comprising of Kindlin-2, Stat3 and Runx2 in articular chondrocytes responsible for maintaining integrity of the articular cartilage and define a potential therapeutic target for OA.

## Introduction

Osteoarthritis (OA) is the most prevalent degenerative joint disease and the leading cause of chronic disability among elderly people worldwide. The etiologic factors of human OA include aging, joint overuse or injury, obesity and heredity. While OA is a whole joint disease affecting articular cartilage, subchondral bone and synovium, a progressive loss of articular cartilage is a hallmark event of OA pathology (*1*). Molecular mechanisms underlying OA initiation, development and progression remain elusive. As a consequence, there are currently no FDA-approved OA treatments or effective interventions to limit OA progression (*2*). Therefore, it is important to define mechanisms that control the articular cartilage homeostasis under physiological condition and how they are altered under OA state.

Chondrocytes are the only cell type in the articular cartilage and are surrounded by a collagen-rich extracellular matrix (ECM), the major target of osteoarthritic cartilage degradation. A hypertrophic and catabolic phenotype characterized by aberrant production of ECM-degrading proteases Mmp13 and Adamts4/5 by articular chondrocytes facilitates ECM degradation and OA initiation and progression (*3, 4*). While integrins are the transmembrane receptors for ECM, whether and how alterations in the ECM-integrin signaling pathway are involved in OA initiation and progression are still controversial (*5–7*). Several studies using genetically modified mouse models reveal that aberrant accumulation of Runx2 protein accelerates articular chondrocyte hypertrophy and stimulates expression of ECM-degrading proteases, leading to OA (*4, 8–14*). Furthermore, activation of the signal transducer and activator of transcription (Stat3) stimulates, while inactivation of Stat3 inhibits, OA initiation and progression (*15–17*). However, key signaling molecules that maintain the articular chondrocyte anabolism and integrity of the articular cartilage remain poorly understood. Furthermore, it is important to investigate whether alterations in expression of these molecules in articular chondrocytes play major roles in pathogenesis of OA.

Kindlins are key focal adhesion proteins that interact with the cytoplasmic domain of the β integrins and activate integrins to regulate cell-ECM adhesion, migration and signaling (*18–20*). In mammalian cells, there are three Kindlin proteins, i.e., Kindlin-1, −2 and −3, encoded by *Fermt1*, *Fermt2* and *Fermt3,* respectively (*21, 22*). Human genetic diseases are linked to mutations in Kindlin-1 and −3, but not Kindlin-2 (*23–26*). Kindlin-2 is essential for early embryonic development; thus, global inactivation of the gene encoding Kindlin-2 resulted in very early embryonic lethality at E7.5 in mice (*27*). Previous studies of Kindlin-2 primarily focus on its roles in regulation of tumor formation, progression and metastasis (*28*). Recently, increasing attention has been paid to its roles in control of organogenesis and homeostasis through both integrin-dependent and integrin-independent mechanisms (*28–40*). We recently demonstrate that Kindlin-2 plays critical roles in regulation of skeletal development and bone remodeling through distinct molecular mechanisms (*41–44*). However, it is not known whether Kindlin-2 has a role in articular cartilage homeostasis and whether alterations in its expression in articular chondrocytes are involved in OA initiation and progression.

In this study, we demonstrate that Kindlin-2, but not Kindlin-1 and −3, is highly expressed in articular chondrocytes of healthy articular cartilage and dramatically down-regulated in the degenerated articular cartilage of aged mice and patients with OA. We demonstrate that Kindlin-2 loss in chondrocytes causes spontaneous OA and exacerbates instability-induced OA in adult mice. Kindlin-2 deficiency promotes hypertrophic differentiation and matrix catabolism through Stat3-dependent up-regulation of Runx2 in articular chondrocytes, leading to OA. Intraarticular injection of AAV5 expressing Kindlin-2 attenuates OA damages caused by aging and instability in mice.

## Results

### Kindlin-2, but not Kindlin-1 or −3, is highly expressed in chondrocytes of the hyaline articular cartilage in mice and humans

As an initial step to investigate potential role of the Kindlin proteins in the articular cartilage, we examined their expression by performing the Safranin O & Fast Green (SO&FG) and immunofluorescence (IF) staining of serial knee joint sections from adult C57BL/6 mice using antibodies against Kindlin-1, −2 or −3 (Figure 1a, top). Results revealed that Kindlin-2 was strongly detected in articular chondrocytes of the hyaline cartilage (Figure 1a,b). In contrast, both Kindlin-1 and −3 proteins were not expressed in the articular chondrocytes. It is interesting to observe that expression of Kindlin-2 was essentially lost in chondrocytes of the calcified cartilage (Figure 1a). Similarly, Kindlin-2, but not Kindlin-1 and 3, was highly expressed in articular chondrocytes of the human knee joint cartilage (Figure 1a,c).

**Figure 1.**
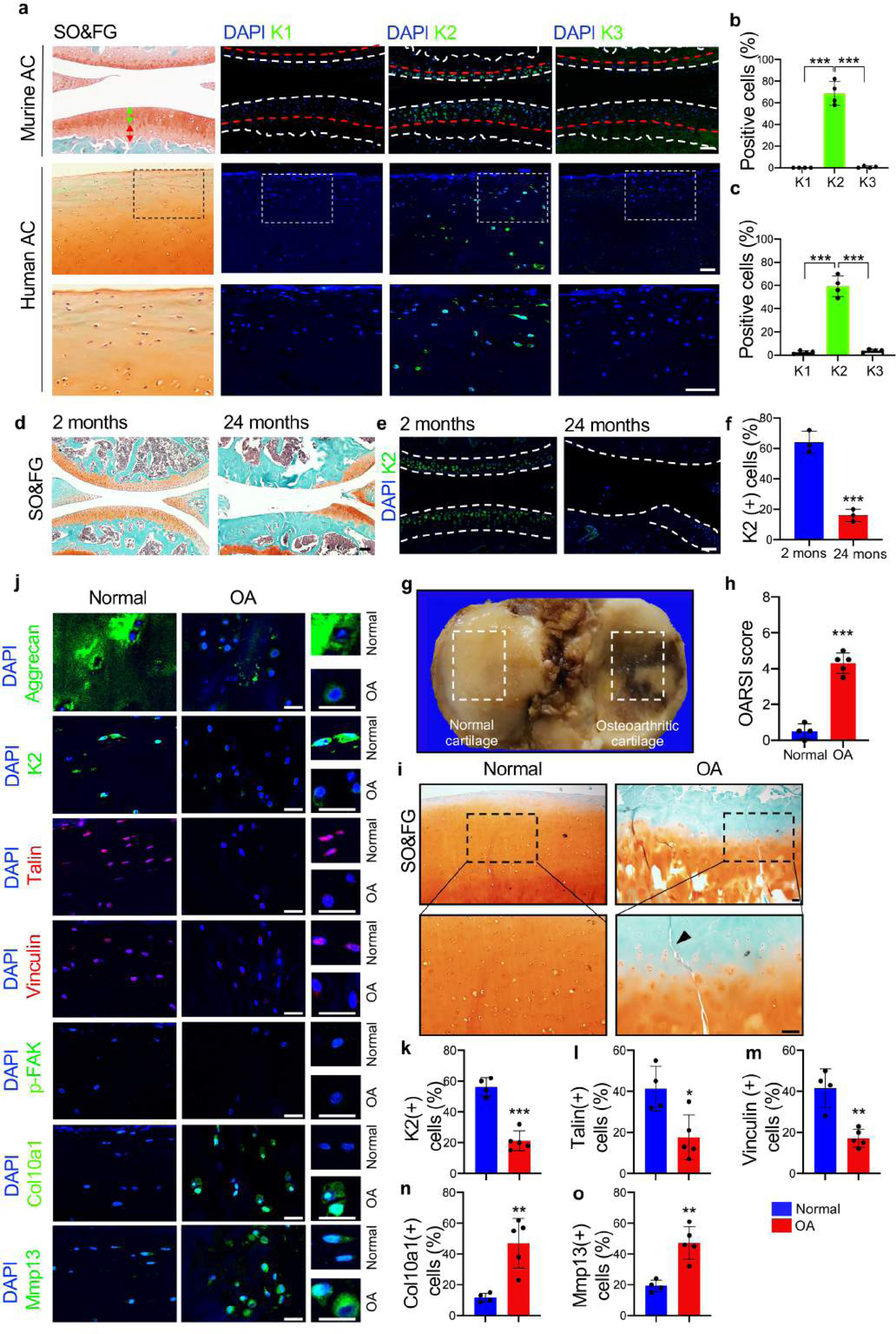
Kindlin-2 is highly expressed in chondrocyte of the hyaline articular cartilage and is reduced in aged mouse and human OA cartilages. (**a-c**) Safranin O & Fast Green (SO&FG) and immunofluorescent (IF) staining of serial sections of mouse (top) and human (bottom) knee joint cartilage. Green double headed arrow indicates hyaline cartilage; Red double headed arrow indicates the calcified cartilage. Red dashed line indicates the tide mark. Scale bar: 50 μm. Quantification of Kindlin1-, 2- and −3-positive cells in articular cartilage (b,c). (**d-f**) SO&FG and IF staining of serial knee joint sections from young (2-mo-old) and aged (24-mo-old) mice. Scale bar: 50 μm. Quantification of Kindlin-2-positive cells in cartilage (f). (**g**) Human knee joint articular cartilages were obtained from total knee replacement of OA patients. White dashed boxes indicate respective normal and osteoarthritic (OA) areas. (**h**) Osteoarthritis Research Society International (OARSI) score of normal and OA cartilages. (**i**) SO&FG staining of normal and OA cartilages. Higher magnification images of dashed boxed areas (bottom panels). Scale bar: 50 μm. Black arrowhead indicates a vertical fissure in OA cartilages. (**j-o**) IF staining of normal and OA cartilage sections for expression of Kindlin-2, talin, vinculin, p-FAK, Col10a1 and Mmp13. Scale bar: 50 μm. Quantitative data (k-o). Results are expressed as mean ± standard deviation (s.d.). \**P* < 0.05, \*\**P* < 0.01, \*\*\**P* < 0.001.

### Kindlin-2 expression is drastically reduced in the degenerated articular cartilage of aged mice and patients with OA

We found that the number of Kindlin-2-positive articular chondrocytes in the knee joints was decreased by 4-fold in aged (24 mo) mice compared to that in young (2 mo) mice (Figure 1d-f) (64% in 2 mo versus 16% in 24 mo, *P* < 0.0005, Student’s *t* test). It should be noted that 24-mo-old mice displayed a dramatic degeneration of the knee joint articular cartilage (Figure 1d). We next obtained human knee joint cartilage samples from total knee arthroplasty (TKA) (Figure 1g). As expected, OA cartilage displayed a dramatic increase in OARSI (Osteoarthritis Research Society International) score (Figure 1h) and a decrease in aggrecan-containing cartilage (Figure 1i). Please note a vertical fissure in OA cartilage (Figure 1i). Results from IF staining showed a drastic reduction in expression of aggrecan in OA versus normal cartilage (Figure 1j). The percentages of Kindlin-2-, talin- and vinculin-positive cells were all dramatically reduced in OA relative to normal cartilage (Figure 1j-m), while those of both Col10a1- and Mmp13-positive cells were significantly increased in OA versus normal cartilage (Figure 1j,n,o). The percentage of Kindlin-2-positive articular chondrocytes was decreased by 2.7-fold in OA versus normal cartilage (Figure 1k) (56.25% in Normal versus 21.2% in OA, *P* < 0.0001, Student’s *t* test). Note: the expression level of p-FAK was extremely low in both OA and normal cartilage (Figure 1j).

### Inducible deletion of Kindlin-2 in chondrocytes at adult stage causes striking spontaneous OA-like phenotype

Based on above observations, we wondered whether the loss of Kindlin-2 in articular chondrocytes plays a role in promotion of OA development and progression. To test if this is the case, we generated mice bearing conditional alleles of *Kindlin-2* and *Aggrecan^CreERT2^*, i.e., *K2^fl/fl^; Aggrecan^CreERT2^* (Supplementary Figure 1). Note: a high Cre-recombination efficiency was observed in the knee joint articular chondrocytes, but not in cells of the synovium, in *Aggrecan^CreERT2^* mice at 4 weeks after tamoxifen injections (Supplementary Figure 2a). At 2 months of age, *K2^fl/fl^; Aggrecan^CreERT2^* mice were subjected to five daily injections of tamoxifen (TM) (100 mg/kg body weight) to generate the chondrocyte conditional Kindlin-2 knockout mice (hereafter referred to as cKO) (Figure 2a). The *K2^fl/fl^; Aggrecan^CreERT2^* mice injected with corn oil were used as controls in this study. At 6 months after TM injection (same hereinafter), we observed a marked enlargement of the knee joint (left panel, red dashed line) (Figure 2b), an excessive tibial plateau angle (middle panel, red double headed arrow) (Figure 2b) and articular cartilage damage of the femoral condyles (right panel, red arrows) in cKO mice (Figure 2b). X-ray micro-computerized tomography (μCT) imaging of the knee joints revealed increasing osteophyte formation in cKO but not in control mice (Figure 2c,d and Supplementary Figure 3a). cKO mice displayed increased volume of calcified meniscus and synovium (Figure 2d) and hyperalgesia (Figure 2e). Kindlin-2 loss caused a dramatic loss of the articular cartilage, as demonstrated by a dramatic increase in OARSI score and a decrease in cartilage area in cKO mice (Figure 2f,i,j and Supplementary Figure 3b-d). Kindlin-2 loss stimulated a synovial hyperplasia (Figure 2g,l). At the molecular level, the numbers of Runx2-, Col10a1- and Mmp13-positive cells were dramatically increased in cKO versus control articular cartilage (Figure 2h,m-p and Supplementary Figure 4a-f). In fact, those cells were rarely observed in control articular cartilage. As expected, Runx2-positive cells were also detected in the calcified cartilage, subchondral bone and bone marrow in both control and cKO mice.

**Figure 2.**
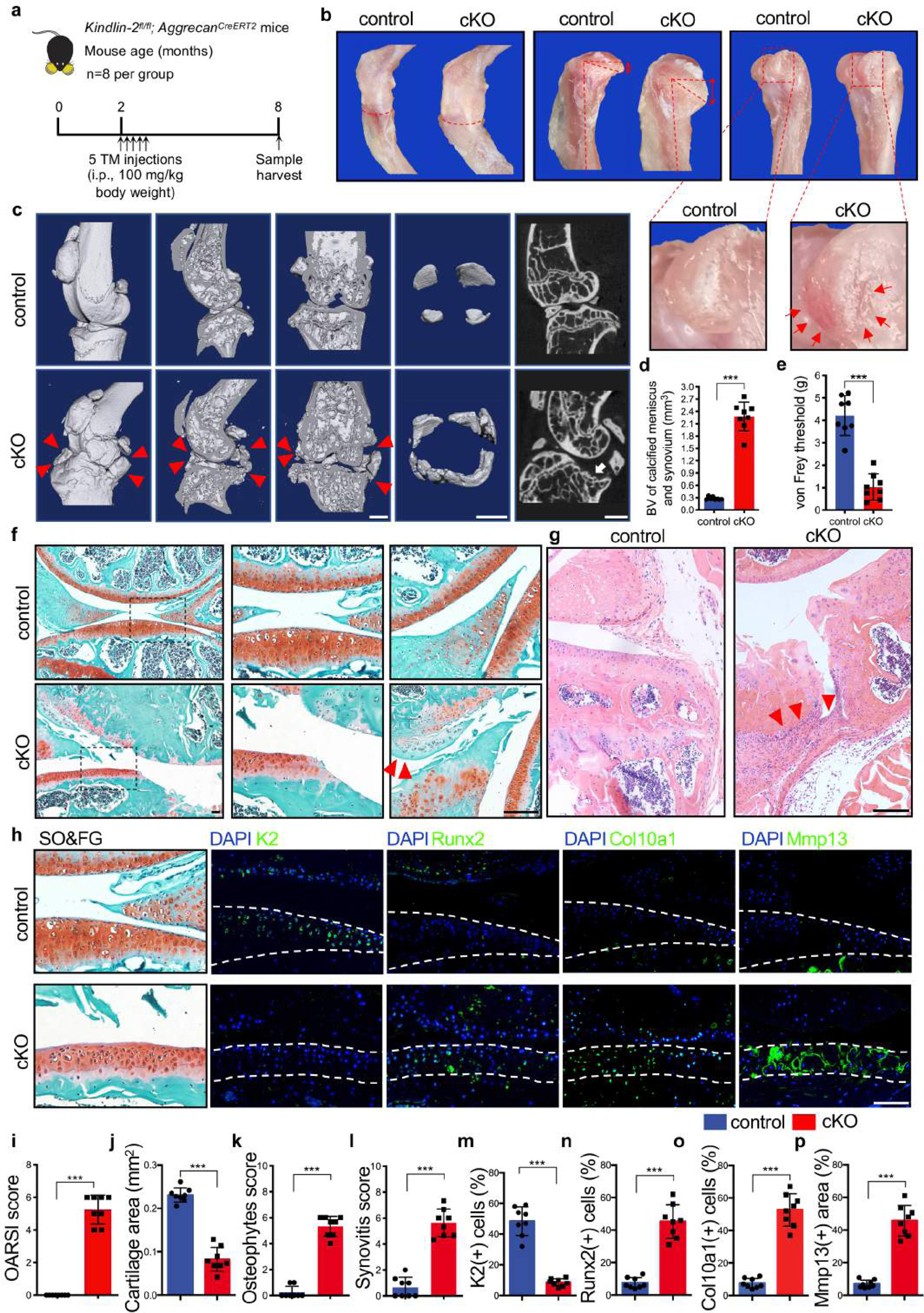
Inducible deletion of Kindlin-2 in chondrocytes causes striking spontaneous OA in adult mice. (**a**) A schematic diagram illustrating the experimental design. At 2 months of age, *Kindlin-2^fl/fl^; Aggrecan^CreERT2^* male mice received five daily intraperitoneal injections of tamoxifen (TM) (cKO, *N* = 8) and corn oil (control, *N* = 8). Six months after TM injection, mice were sacrificed, and knee joints were collected. (**b**) Representative images showing enlargement of the knee joint (left panel, red dashed line), excessive tibial plateau angle (middle panel, red double headed arrow) and cartilage damage of femoral condyles (right panel, red arrow) in cKO mice. Higher magnification images of dashed boxed areas (lower panels). (**c**) Three-dimensional (3D) reconstruction from micro-computerized tomography (μCT) scans of control and cKO knee joints. Scale bar, 1.0 mm. (**d**) The volume of calcified meniscus and synovial tissue was analyzed by μCT. (**e**) von Frey test. Three months after TM injection, cKO male mice display a hyperalgesia with a dramatic reduction in the 50% paw withdrawal threshold. (**f**) SO&FG staining of control and cKO knee joint sections (left panel). Higher magnification images showing dramatic articular cartilage loss (middle panels) and osteophyte outgrowth (right panels, red arrowheads). Scale bar: 50 μm. (**g**) H&E staining of control and cKO knee joint sections. Arrowheads show marked synovial hyperplasia. Scale bar: 50 μm. (**h**) SO&FG and IF staining of serial knee joint sections were performed to determine expression of Kindlin-2, Runx2, Col10a1 and Mmp13 in articular cartilage. Scale bar: 50μm. (**i**-**l**) Quantification of OARSI score (i), cartilage area (j), osteophyte score (k) and synovitis score (l) was performed using histological sections. (**m**-**p**) Quantitative data of expression of Kindlin-2 (m), Runx2 (n), Cola10a1 (o) and Mmp13 (p). Results are expressed as mean ± standard deviation (s.d.). \*\*\**P* < 0.001.

Collectively, we demonstrate that Kindlin-2 loss in adult mice promotes expression of Runx2, chondrocyte hypertrophic and catabolic phenotype, and spontaneous OA-like phenotypes, including progressive cartilage loss and structural deterioration, osteophyte outgrowth, synovial hyperplasia and pain, which mimic major pathological features of human OA.

### Kindlin-2 deficiency in chondrocytes at adult stage exacerbates instability-induced OA

We further investigated whether Kindlin-2 loss impacts progression of the injury-induced OA by utilizing a well-established destabilization of the medial meniscus (DMM) mouse OA model. At 2 months of age, the *K2^fl/fl^; Aggrecan^CreERT2^* mice were subjected to sham or DMM surgery. One week later, mice were injected with TM (cKO) or corn oil (control) as indicated in Figure 3a. At 8 weeks after surgery, we performed μCT scans and histomorphometrical analyses of SO&FG and H/E-stained knee joint sections to evaluate knee joint damages (Figure 3b-e). At this time point, cKO mice did not display marked abnormalities in the volume of calcified meniscus and synovial tissue (Figure 3c), OARSI score (Figure 3f), cartilage area (Figure 3g), osteophyte score (Figure 3h) and synovitis score (Figure 3i). As expected, when compared to control mice with sham operation, control mice with DMM (control-DMM) exhibited apparent OA lesions, as revealed by a dramatic loss of the articular cartilage, synovial hyperplasia and osteophyte outgrowth (Figure 3b-i) (*P* < 0.05, control-sham vs control-DMM for all indicated parameters). Importantly, cKO mice with DMM (cKO-DMM) displayed more severe OA phenotypes than control-DMM did (Figure 3b-i) (*P* < 0.05, control-DMM vs cKO-DMM for all indicated parameters). Collectively, Kindlin-2 deletion in chondrocytes exacerbates OA lesions caused by instability in mice.

**Figure 3.**
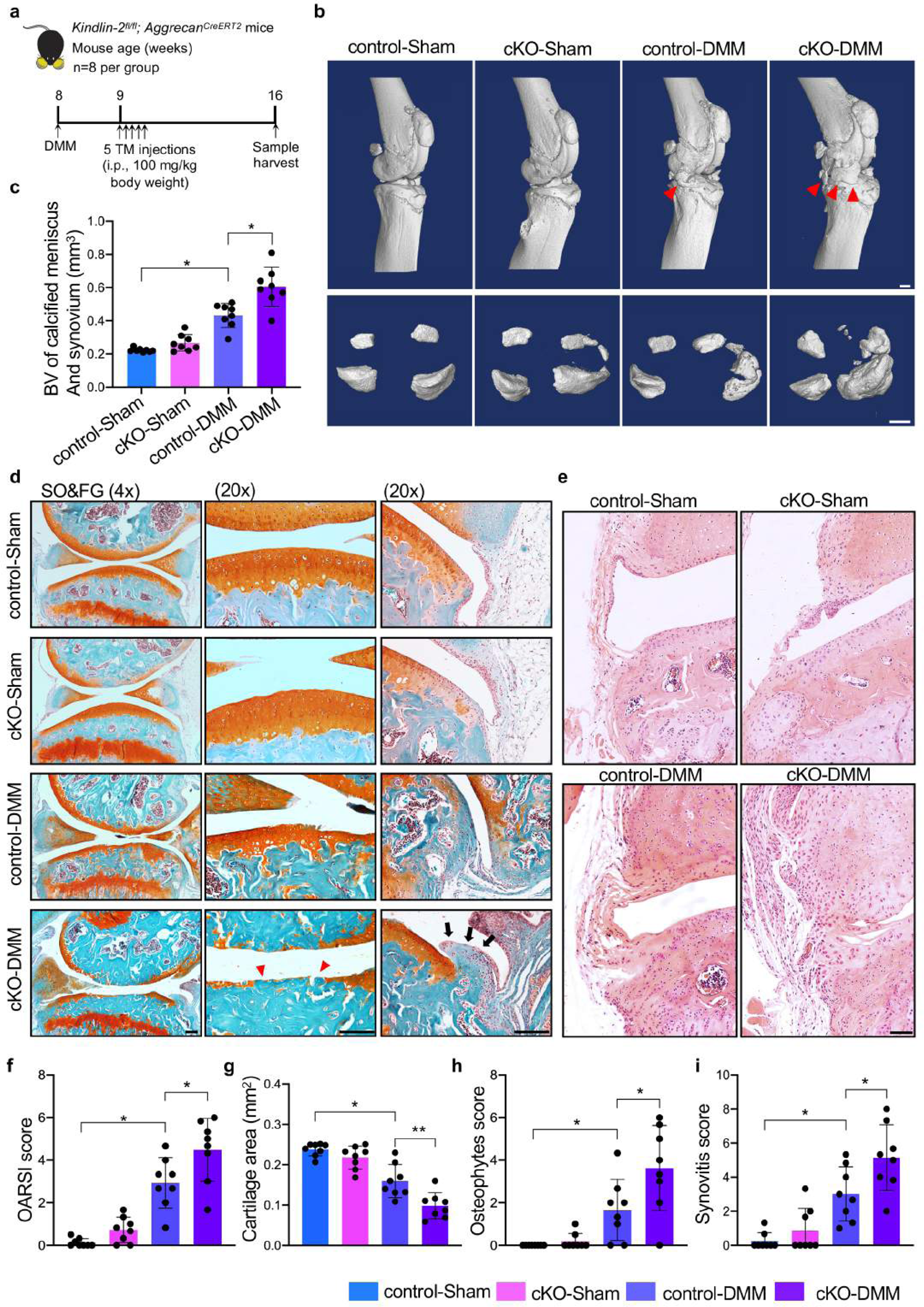
Kindlin-2 deficiency in chondrocytes accelerates OA progression in mice with DMM surgery. (**a**) A schematic diagram illustrating the experimental design. (**b**) μCT scans of knee joints from control and cKO mice at 8 weeks after sham or DMM surgery. *N* = 8 per group. Scale bar, 1.00 mm. (**c**) The volume of calcified meniscus and synovial tissue was analyzed by μCT. (**d**) Representative images of SO&FG-stained sections of control and cKO knee joints. Higher magnification images (right two panels) showing exacerbated cartilage loss (red arrowheads) and osteophyte outgrowth (black arrows) in cKO/DMM group. Scale bar: 50 μm. (**e**) Representative images of H&E staining. Scale bar: 50 μm. (**f-i**) Quantification of OARSI score (f), cartilage area (g), osteophyte score (h) and synovitis score (i) were performed using histological sections. Results are expressed as mean ± standard deviation (s.d.). *\*P* < 0.05.

### Kindlin-2 loss up-regulates expression of p-Sta3, Runx2, Col10a1 and ECM-degrading proteases in articular chondrocytes

We performed RNA sequencing analysis using total RNA from the knee joints of control and cKO mice at 5 months after TM injections. Results from KEGG pathway analysis revealed significant enrichment of several signaling pathways, including the JAK-STAT3 and NF-κB signaling pathways (Figure 4a). Consistent with results from the RNAseq analysis, knockdown of Kindlin-2 increased the protein levels of p-Stat3 (but not its total protein, t-Stat3) in ATDC5 cells and in primary articular chondrocytes (Figure 4b,c and Supplementary Figures 5, 6). Furthermore, Kindlin-2 loss increased the protein levels of Runx2, Col10a1, Mmp13 and Adamts5 in ATDC5 cells and in primary articular chondrocytes (Figure 4b,c and Supplementary Figures 5, 6). IF staining showed that Kindlin-2 knockdown increased expression of p-Stat3 in nuclei of ATDC5 cells and that of p-Stat3 and Runx2 in primary articular chondrocytes (Figure 4d and Supplementary Figure 6e). Furthermore, Kindlin-2 knockdown increased the protein level of p-Stat3 (Y705), but not p-Stat3 (Y727), in ATDC5 cells (Figure 4e and Supplementary Figure 7a). Results from IF staining of serial sections of the knee joints revealed that percentage of p-Stat3-positive chondrocytes was drastically increased in cKO versus control cartilage (Figure 4f,g). Furthermore, the percentage of activated β1 integrin-positive cells was significantly reduced, whereas the percentages of p-p38-, p-Erk- and p-Jak2-positive cells were not significantly altered in cKO articular cartilages compared to those in control articular cartilages (Supplementary Figure 8a-e). Loss of Kindlin-2 markedly impaired the attachment and spreading of primary articular chondrocytes on collagen-II coated surface in vitro (Supplementary Figure 9).

**Figure 4:**
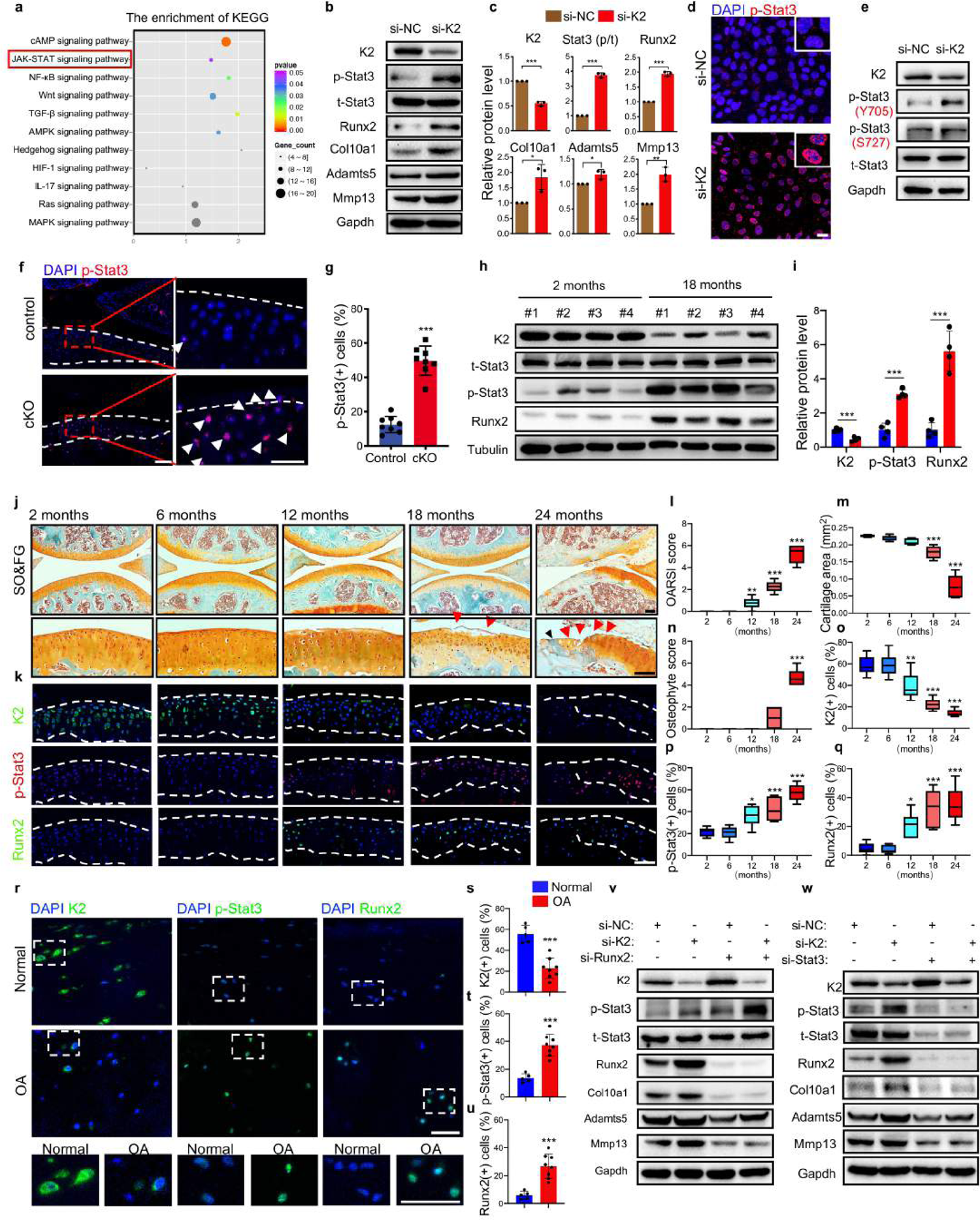
Kindlin-2 loss induces chondrocyte hypertrophic differentiation and catabolism through Stat3-dependent up-regulation of Runx2. (**a**) KEGG pathway analysis of cellular signaling pathways enriched in RNA-seq analysis using RNAs from of control and cKO articular cartilages (3 mice per group) at 5 months after TM injections. (**b**) ATDC5 cells were transfected with si-NC and si-K2 for 48h, followed by western blotting for expression of the indicated proteins. (**c**) Quantification of (b). Experiments were repeated three times independently. (**d**) IF staining of p-Stat3 in ATDC5 cells transfected with si-NC and si-K2. Scale bar: 25 μm. (**e**) ATDC5 cells were transfected with si-NC and si-K2 for 48h, followed by western blotting for expression of the indicated proteins. (**f**) IF staining for expression of p-Stat3 in articular cartilage of control and cKO mice at 3 months after TM induction. Higher magnification images of red dashed boxed areas (right panels). White arrowheads indicate the elevated expression of p-Stat3 in articular chondrocytes. Scale bar: 50 μm. (**g**) Quantification of (f). (**h**) Western blotting analyses of expression of Kindlin-2, t-Stat3, p-Stat3 and Runx2 in articular cartilages from young (2-mo) and aged (18-mo) C57BL/6 mice. (**i**) Quantification of (h). (**j**) Representative images of SO&FG-stained sections of knee joints of C57BL/6 mice at different ages. Higher magnification images (lower panel) showing dramatic cartilage loss (red arrowheads) and osteophyte outgrowth (black arrowhead) in 18- and 24-mo-old mice. Scale bar: 50 μm. (**k**) IF staining for expression of Kindlin-2, p-Stat3 and Runx2 in articular cartilage of C57BL/6 mice at different ages. (**l-q**) Quantification of OARSI score (l), cartilage area (m), osteophyte score (n), Kindlin-2 (o), p-Stat3 (p) and Runx2 (q) in articular cartilages from C57BL/6 mice during aging. *N* = 6 per group. (**r-u**) IF staining for expression of Kindlin-2, p-Stat3 and Runx2 in human normal and OA articular cartilages. Scale bar: 50 μm. Quantitative data (s-u). *N* = 5 for normal, *N* = 8 for OA. (**v**) Runx2 knockdown. ATDC5 cells were transfected with si-NC or si-K2 with and without si-Runx2, followed by western blotting. (**w**) Stat3 knockdown. ATDC5 cells were transfected with si-NC or si-K2 with and without si-Stat3, followed by western blotting. All data are expressed as mean ± standard deviation (s.d.) *\*P* < 0.05, *\*\*P* < 0.01, *\*\*\*P* < 0.001.

**Figure 5.**
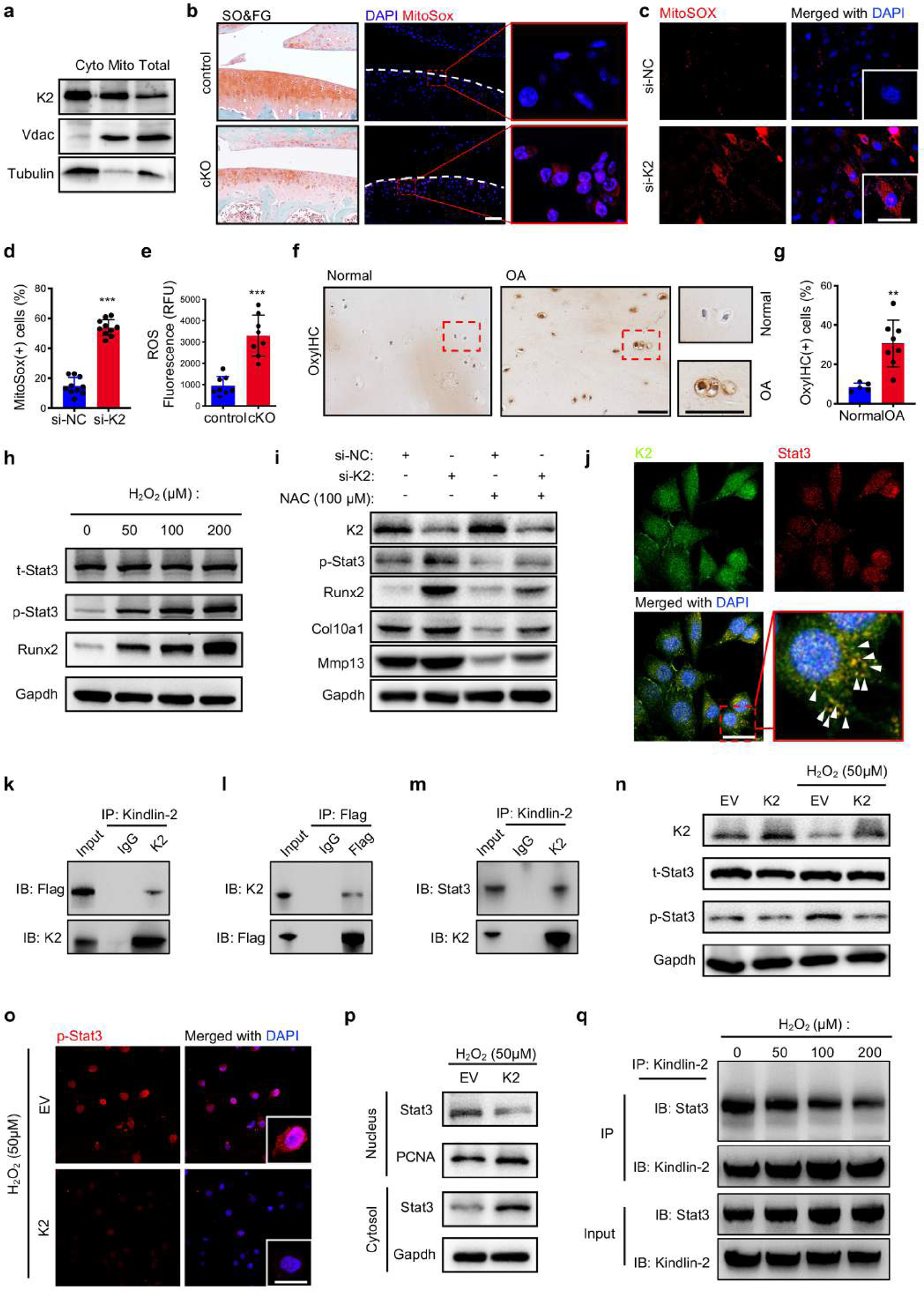
Kindlin-2 deficiency promotes oxidative stress, Stat3 activation and nuclear translocation in chondrocytes. (**a**) Primary articular chondrocytes were isolated from 2-month-old *Kindlin-2^fl/fl^; Aggrecan^CreERT2^* male mice. The cytosolic fraction (Cyto, left lane), mitochondrial fraction (Mito, middle lane) and total cell lysates (Total, right lane) of the primary chondrocytes were isolated and analyzed by western blotting with antibodies against Kindlin-2, Vdac (voltage-dependent anion channel, a mitochondrial marker protein) and tubulin. (**b**) The mitochondrial ROS levels in articular cartilages of control and cKO mice at 3 months after TM injections were visualized by MitoSOX red staining. Scale bar: 50 μm. (**c,d**) Increased level of mitochondrial ROS in ATDC5 cells transfected with control (si-NC) and Kindlin-2 siRNA (si-K2) for 48h. Scale bar: 25 μm. Data were quantified with 10 independent fields of view and shown in (d). (**e)** Elevated ROS levels in articular cartilage extracts isolated from control and cKO mice at 2 months after TM induction (*N* = 8 for each group). (**f**,**g**) OxyIHC staining determining the level of oxidative stress in human normal and OA articular cartilage. Quantitative data were shown in (g). *N* = 5 for normal cartilages, *N* = 8 for OA cartilages. (**h**) Primary articular chondrocytes were treated with the indicated concentrations of H_2_O_2_, followed by western blotting for expression of total Stat3 (t-Stat3), phosphorylated Stat3 (p-Stat3) and Runx2. (**i**) ATDC5 cells were transfected with si-NC or si-K2 with and without NAC (100 μM), followed by western blotting. (**j**) Co-localization of Kindlin-2 and Stat3 in primary articular chondrocytes. Scale bar, 25 μm. (**k**-**l**) Co-immunoprecipitation (co-IP) assay. COS-7 cells were co-transfected with plasmids expressing Flag-Stat3 and full-length Kindlin-2. Protein extracts were incubated with either Kindlin-2 antibody (k) or Flag antibody (l), followed by western blotting using Flag and Kindlin-2 antibodies. (**m**) co-IP assay. Protein extracts from primary articular chondrocytes were incubated with Kindlin-2 antibody or IgG, followed by western blotting with antibodies against Stat3 and Kindlin-2. (**n**) ATDC5 cells were transfected with empty vector (EV) and Kindlin-2 expression vector (K2). 24h later, cells were treated with and without 50 μM H_2_O_2_ for another 12h, followed by western blotting using the indicated antibodies. (**o**) IF staining. ATDC5 cells transfected with empty vector (EV) and Kindlin-2 expression vector (K2). 24h later, cells were treated with and without 50 μM H_2_O_2_ for another 12h, followed by IF staining with p-Stat3 antibody and DAPI. Scale bar, 25 μm. (**p**) Nuclear translocation of Stat3 and K2 overexpression. ATDC5 cells transfected with empty vector (EV) and Kindlin-2 expression vector (K2). 24h later, cells were treated with and without 50 μM H_2_O_2_ for another 12h. Nuclear proteins and cytoplasmic proteins were separated and the expression patterns of Stat3 protein in nucleus and cytosol were detected by western blotting. PCNA (proliferating cell nuclear antigen) and Gapdh were used as control for nuclear and cytoplasmic proteins, respectively. (**q**) co-IP assay. Primary articular chondrocytes were treated with increasing concentrations of H_2_O_2_ for 12h and protein extracts were incubated with Kindlin-2 antibody, followed by western blotting with antibodies against Stat3 and Kindlin-2.

**Figure 6.**
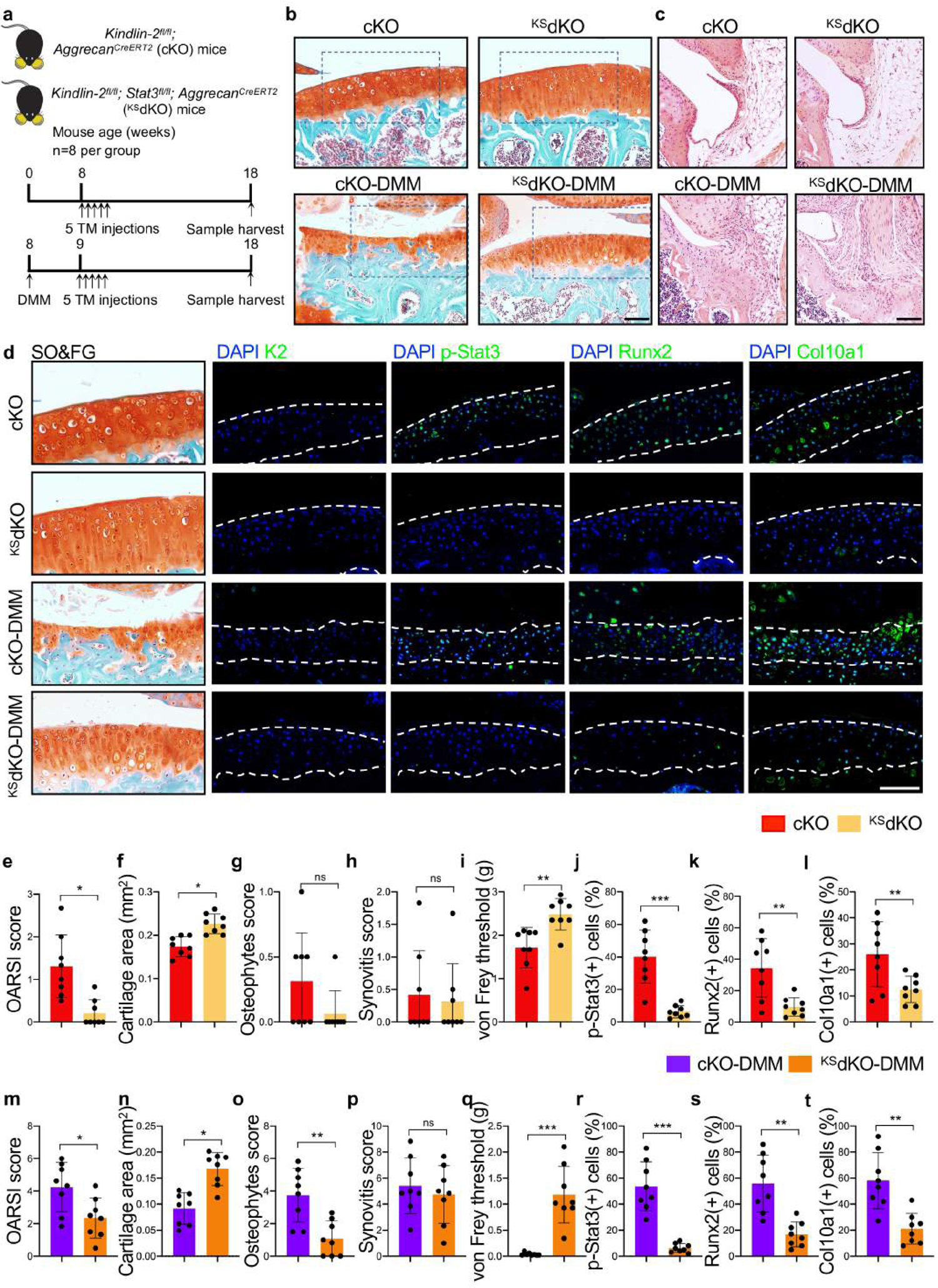
Genetic deletion of Stat3 in chondrocyte corrects Runx2 accumulation and attenuates OA lesions in cKO mice. (**a**) A schematic diagram illustrating the experimental design. (**b**) Representative images of SO&FG-stained sections of knee joints from cKO and ^KS^dKO mice at 10 weeks after TM injection (upper panel) or at 8 weeks after DMM (lower panel). *N* = 8 per group. Scale bar, 50 μm. (**c**) Representative images of H&E staining. Scale bar: 50 μm. (**d**) SO&FG and IF staining of serial sections of knee joints for expression of Kindlin-2, p-Stat3, Runx2 and Col10a1 in articular cartilage of cKO and ^KS^dKO mice. The white dashed lines indicate the articular cartilage areas. Scale bar: 50 μm. (**e-i,m-q**) Quantitative data of OARSI score (e,m), articular cartilage area (f,n), osteophyte score (g,o), synovitis score (h,p) and von Frey test (i,q). (**j-l,r-t**) Quantitative data for expression of p-Stat3 (j,r), Runx2 (k,s) and Col10a1 (l,t) in articular cartilages of cKO and ^KS^dKO mice with or without DMM. All data are expressed as mean ± standard deviation (s.d.). \**P* < 0.05, \*\**P* < 0.01, \*\*\**P* < 0.001.

### Loss of Kindlin-2 expression associates with elevations of p-Stat3 and Runx2 in articular chondrocytes during aging and OA development

Western blotting using protein extracts from articular cartilage of mice with different ages showed that expression of Kindlin-2 was reduced and that of p-Stat3 and Runx2 was increased in aged (18 mo) versus young (2 mo) mice (Figure 4h,i). SO&FG and IF staining of serial knee joints sections of mice with different ages showed that the percentages of Kindlin-2-positive chondrocytes were decreased and those of p-Stat3- and Runx2-positive chondrocytes were increased in aged versus young cartilages (Figure 4j,k). Quantitative data showed that the severity of aging-induced OA, as measured by increases in OARSI score and osteophyte score and decrease in cartilage area, was closely correlated to the magnitude of Kindlin-2 down-regulation or p-Stat3 and Runx2 up-regulation in articular chondrocytes (Figure 4l-q). In humans, the percentages of p-Stat3- and Runx2-positive chondrocytes were increased by 2.7- and 4.5-fold, respectively, in OA relative to normal cartilage (*P* < 0.0001, normal versus OA, Student’s *t* test) (Figure 4r,t,u). Again, the percentage of Kindlin-2-positive chondrocytes was decreased in human OA versus normal cartilage (Figure 4r,s).

### Kindlin-2 deficiency stimulates chondrocyte hypertrophic differentiation and catabolism through Stat3-dependent up-regulation of Runx2 expression

Runx2 is known to activate expression of Col10a1 and Mmp13, which plays a pivotal role in ECM calcification and OA initiation and progression (*3, 4, 8, 9*). Interestingly, siRNA knockdown of Runx2 essentially abolished Kindlin-2 loss-stimulated up-regulation of Col10a1, Mmp13 and Adamts5 without down-regulating expression of p-Stat3 (Figure 4v and Supplementary Figure 7b,d). A previous in vitro study showed that Stat3 transcriptionally activates expression of Runx2 to promote human osteoblastic differentiation (*45*). Based on these observations, we wondered whether Kindlin-2 loss stimulates the chondrocyte hypertrophic and catabolic phenotypes by up-regulating Runx2 through Stat3. In support of this notion, we found that siRNA knockdown of Stat3 largely reversed the Runx2 mRNA and protein accumulation caused by Kindlin-2 loss (Figure 4w and Supplementary Figures 7c,e,f). Furthermore, Stat3 knockdown essentially abolished the up-regulation of Col10a1, Mmp13 and Adamts5 caused by Kindlin-2 loss (Figure 4w and Supplementary Figure 7c,e,f). Stat3 siRNA decreased, while Stat3 overexpression increased, the level of Runx2 protein in ATDC5 cells (Supplementary Figure 7g,h). Collectively, these results suggest that Kindlin-2 deficiency induces chondrocyte hypertrophic differentiation and catabolism by Stat3-dependent up-regulation of Runx2 in chondrocytes.

**Figure 7.**
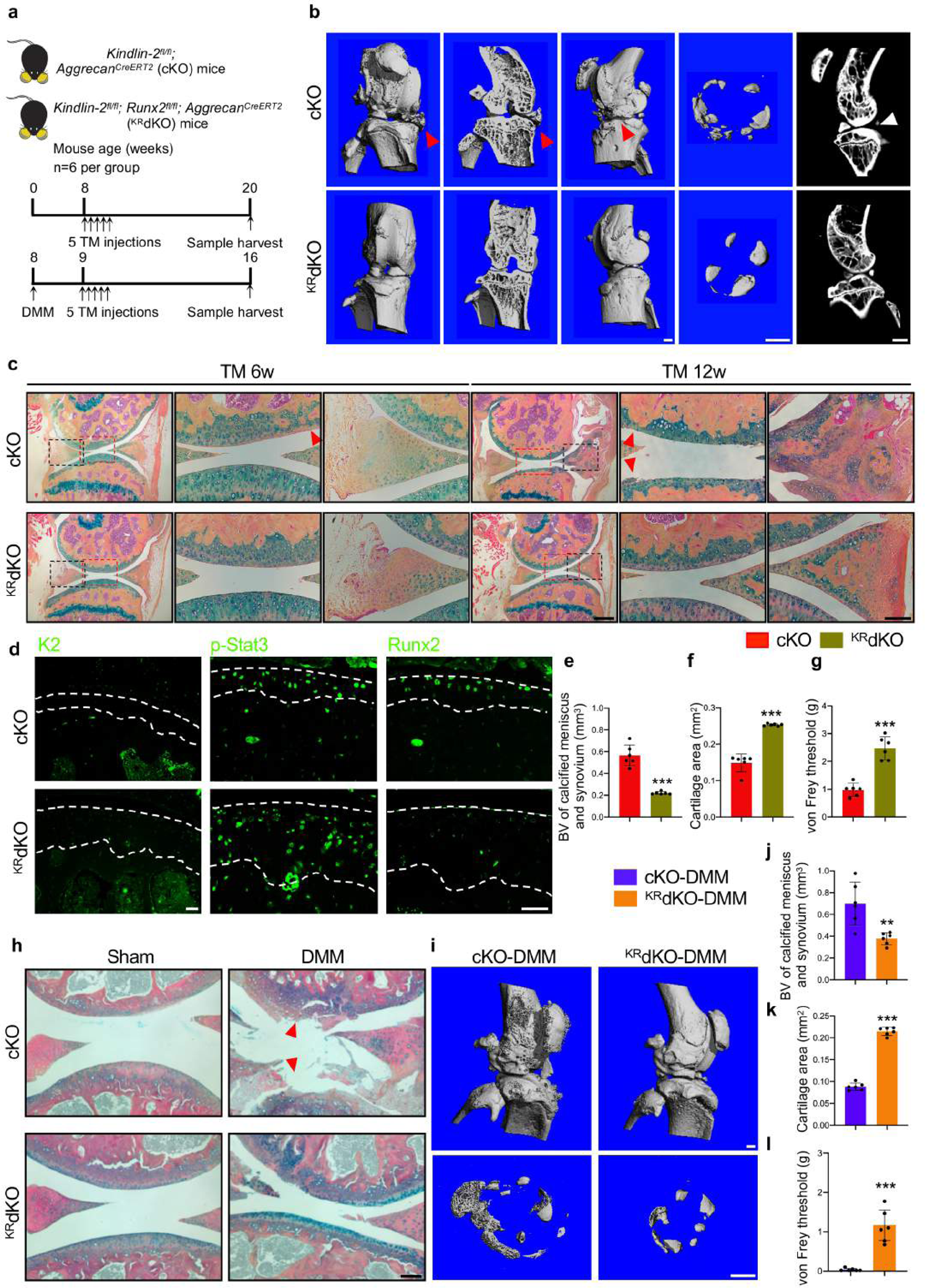
Genetical deletion of Runx2 in chondrocytes palliates spontaneous and DMM-induced OA defects in cKO mice. (**a**) A schematic diagram illustrating the experimental design. (**b**) μCT scans of knee joints from cKO and ^KR^dKO mice at 12 weeks after TM injection. Scale bar, 1 mm. (**c**) Representative images of Alcian Blue and Orange G (AB&OG)-stained sections of knee joints from cKO and ^KR^dKO mice at 6 weeks (left 3 panels) or at 12 weeks (right 3 panels) after TM injection. Scale bar, 50 μm. (**d**) IF staining for expression of Kindlin-2, p-Stat3 and Runx2 in cKO and ^KR^dKO articular cartilage. The white dashed lines indicate the articular cartilage areas. Scale bar: 50 μm. (**e-g**) Quantitative data of the volume of calcified meniscus and synovial tissue (e), cartilage area (f) and von Frey test (g) at 12 weeks after TM injections. *N* = 6 mice per group. (**h**) Representative images of AB&OG-stained sections of knee joints from cKO and ^KR^dKO mice at 8 weeks after DMM. Red arrowheads indicate articular cartilage destruction. Scale bar, 50 μm. (**i**) μCT scans of knee joints from cKO and ^KR^dKO mice at 8 weeks after DMM. Scale bar, 1 mm. (**j-l**) Quantitative data of the volume of calcified meniscus and synovial tissue (j), cartilage area (k) and von Frey test (l) at 8 weeks after DMM. *N* = 6 mice per group. All data are expressed as mean ± standard deviation (s.d.). \**P* < 0.05.

### Kindlin-2 loss accumulates reactive oxygen species (ROS) to activate Stat3 in chondrocytes

We recently reported that a fraction of the Kindlin-2 was present in mitochondrion of lung cancer cells (*34*). Consistent with this finding, Kindlin-2 was also detected in extracts of the mitochondrial fraction of primary articular chondrocytes (Figure 5a). This result was specific because the voltage-dependent anion channel (Vadc), a mitochondrial marker, was detected in mitochondrial but not cytoplasmic fraction of chondrocytes (Figure 5a). Since mitochondrion is the primary source for the intracellular reactive oxygen species (ROS) (*46*), we determined whether Kindlin-2 loss increases ROS production in chondrocytes. Strikingly, we found that the level of mitochondrial ROS, as visualized by the MitoSOX red staining of the knee joint sections, was dramatically elevated in articular chondrocytes or cartilage extracts from the knee joint of cKO mice at 3 months after TM injections (Figure 5b,e). Furthermore, siRNA knockdown of Kindlin-2 increased the level of mitochondrial ROS in ATDC5 cells (Figure 5c,d). Interestingly, excessive oxidative stress in articular chondrocytes was observed in the damaged knee joint articular cartilage from human OA patients, as revealed as OxyIHC staining (Figure 5f,g). H_2_O_2_, a common ROS in the cells, dose-dependently increased the levels of p-Stat3 and Runx2 proteins without affecting expression of t-Stat3 protein in primary articular chondrocytes (Figure 5h). Furthermore, up-regulations of p-Stat3, Runx2, Col10a1 and Mmp13 caused by Kindlin-2 knockdown were largely reversed by N-acetyl cysteine (NAC) (Figure 5i), a potent ROS scavenger.

### Kindlin-2 interacts with Stat3 and inhibits Stat3 nuclear translocation in chondrocytes

To further explore mechanisms through which Kindlin-2 deficiency activates Stat3, we determined whether Kindlin-2 interacts with Stat3 by performing immunofluorescence (IF) staining and observed a strong colocalization of both factors in the cytoplasm of primary articular chondrocytes (Figure 5j). We further conducted co-immunoprecipitation (co-IP) assays using whole cell extracts isolated from the COS-7 cells overexpressing Flag-Stat3 and Kindlin-2 and found that Stat3 was present in the Kindlin-2 immunoprecipitates (Figure 5k) and, vice versa, that Kindlin-2 was present in the Stat3 immunoprecipitates (Figure 5l). Endogenous Kindlin-2 and Stat3 interacted in primary articular chondrocytes (Figure 5m). H_2_O_2_-induced increase in p-Stat3 was abolished by overexpression of Kindlin-2 in ATDC5 cells (Figure 5n,o). Overexpression of Kindlin-2 decreased Stat3 nuclear translocation in ATDC5 cells stimulated by H_2_O_2_ (Figure 5p). Finally, H_2_O_2_ dose-dependently inhibited the Kindlin-2-Stat3 interaction in primary articular chondrocytes (Figure 5q).

### Systemic pharmacological blockade of Stat3 activation palliates cartilage degeneration and osteophyte formation caused by Kindlin-2 loss in mice

We next determined whether Stat3 activation plays a role in Kindlin-2 loss induction of OA by investigating whether systemic inhibition of Stat3 activation by Stattic can mitigate the OA lesions caused by Kindlin-2 loss in mice. In this experiment, at 8 weeks of age, *K2^fl/fl^; Aggrecan^CreERT2^* mice were performed with DMM surgeries. One week later, mice were subjected to TM injections and administration of Stattic through gavage as implicated in Supplementary Figure 10a. Eight weeks later, mice with DMM displayed marked restriction of movement of the hind limb, which was improved by Stattic (Supplementary Figure 10b). Expression of p-Stat3 was strongly detected in cKO mice treated with PBS, which was essentially abolished by Stattic (Supplementary Figure 10c). Furthermore, cartilage loss and osteophyte formation caused by Kindlin-2 deletion were attenuated by Stattic treatment (Supplementary Figure 10c-f). However, synovial hyperplasia in cKO mice was not improved by Stattic (Supplementary Figure 10c,g).

### Genetic deletion of Stat3 in chondrocytes reverses aberrant Runx2 accumulation and ameliorates OA lesions caused by Kindlin-2 deficiency in mice

To obtain further in vivo evidence that Kindlin-2 deletion causes OA by activation of Stat3, we deleted Stat3 expression in chondrocytes and determined its effects on Runx2 expression and OA lesions caused by Kindlin-2 deletion in mice with and without DMM. We crossed the *K2^fl/fl^; Aggrecan^CreERT2^* (cKO) mice with floxed Stat3 mice (*Stat3^fl/fl^*) and generated *K2^fl/fl^; Stat3^fl/fl^; Aggrecan^CreERT2^* mice (hereinafter referred to as ^KS^dKO). At 8 weeks of age, cKO and ^KS^dKO mice were subjected to five TM injections as indicated in Figure 6a (top). Mice were killed at 18 weeks of age. Separately, 8-week-old cKO and ^KS^dKO mice were subjected to DMM surgery, followed by TM treatment as indicated in Figure 6a (bottom). Mice were sacrificed at 16 weeks of age. Results revealed that deletion of Stat3 in chondrocyte corrected the increased OARSI score, osteophyte score and pain and decreased articular cartilage area caused by Kindlin-2 deletion in mice with and without DMM (Figure 6b,c,e-g,i m-o,q). Consistent with results from above Stattic inhibition experiment, Stat3 deletion did not reverse the synovitis stimulated by Kindlin-2 deletion in mice with and without DMM (Figure 6c,h,p). At the molecular level, Stat3 deletion essentially abolished expression of both Runx2 and Col10a1 in articular chondrocytes (Figure 6d,j-l,r-t). As expected, the number of Stat3-positive cells was dramatically decreased in articular chondrocytes in ^KS^dKO mice with and without DMM (Figure 6j,r). Note: deleting one allele of *Stat3* gene in chondrocytes slightly but significantly reversed the cartilage loss caused by Kindlin-2 deletion in cKO mice (Supplementary Figure 12a-c). Collectively, these results support our hypothesis that Stat3 activation plays a critical role in mediation of Kindlin-2 loss-induced Runx2 up-regulation in chondrocytes and OA lesions.

### Deleting Runx2 in chondrocytes reverses OA defects without reducing p-Stat3 expression caused by Kindlin-2 deletion in mice

The next question we asked was whether Runx2 is a major downstream effector of Stat3 in mediation of Kindlin-2 loss-caused OA lesions. We determined whether genetic ablation of Runx2 in chondrocytes can limit the OA lesions caused by Kindlin-2 deletion in mice. We bred the *K2^fl/fl^; Aggrecan^CreERT2^* (cKO) with floxed Runx2 mice (*Runx2^fl/fl^*) and generated *K2^fl/fl^; Runx2 ^fl/fl^; Aggrecan^CreERT2^* mice (hereinafter referred to as ^KR^dKO). We performed two separate sets of experiments on these mice. In the first set of experiment, at 8 weeks of age, cKO and ^KR^dKO mice were treated with TM as indicated in Figure 7a. At 6 and 12 weeks after TM injection, mice were killed, followed by IF staining and histological and μCT analyses of the knee joints. At 12 weeks, we observed significant osteophyte outgrowth in cKO mice, which was essentially abolished in ^KR^dKO mice (Figure 7b,e). A dramatic cartilage loss was observed in cKO mice at 12 weeks, but not at 6-weeks (Figure 7c). The cartilage degeneration and pain caused by Kindlin-2 deficiency were largely attenuated by Runx2 deletion (Figure 7c,f,g). It is important to note that expression of p-Stat3 in chondrocytes was not reduced in ^KR^dKO mice (Figure 7d). As expected, expression of both Kindlin-2 and Runx2 was essentially abolished in ^KR^dKO articular chondrocytes (Figure 7d).

In the second set of experiment, cKO and ^KR^dKO mice were subjected to DMM surgery and TM injections as indicated in Figure 7a. At 8 weeks after DMM surgery, cKO mice displayed marked OA lesions with dramatic cartilage loss, osteophyte formation and pain. These OA lesions were largely reversed in ^KR^dKO mice (Figure 7h-l). Furthermore, the structural deterioration of the knee joint caused by Kindlin-2 deficiency was largely ameliorated in ^KR^dKO mice (Figure 7h,i). It should be noted that the cartilage loss caused by Kindlin-2 deletion in cKO mice was partially reversed by Runx2 haploinsufficiency in chondrocytes (Supplementary Figure 13a-c).

### Intraarticular injection of Kindlin-2-expressing adeno-associated virus decelerates progression of DMM- and aging-induced OA in mice

We next determined whether overexpression of Kindlin-2 via intraarticular injection of Kindlin-2-expressing adeno-associated virus 5 (AAV5) protects against OA development and progression caused by DMM or aging in mice as indicated in Figure 8a. Efficiency of AAV infection in articular chondrocytes was assessed by intraarticular injection of AAV5 expressing enhanced green fluorescent protein (EGFP). Three weeks after injection, strong GFP signal was detected in articular chondrocytes (Figure 8b). Kindlin-2 expressing AAV (AAV5-K2) (5 x 10^9^ particles in 10 μl) or control AAV (AAV5-Con) was intraarticularly injected as we previously described (*11*). Results showed that intraarticular injection of AAV5-K2 markedly increased expression of Kindlin-2 in articular chondrocytes of mice with and without DMM (Figure 8c,e). Importantly, DMM caused dramatic cartilage loss, osteophyte formation, synovial hyperplasia and pain, which were largely ameliorated by AAV5-K2 injection as compared with AVV5-Con group (Figure 8d, f-j). More importantly, aging-induced OA lesions were markedly protected by intraarticular injection of AAV5-K2 (Figure 8k-o). Taken together, these results suggest that targeted expression of Kindlin-2 in articular chondrocytes preserves integrity of articular cartilage and protect against aging- and instability-induced OA.

**Figure 8.**
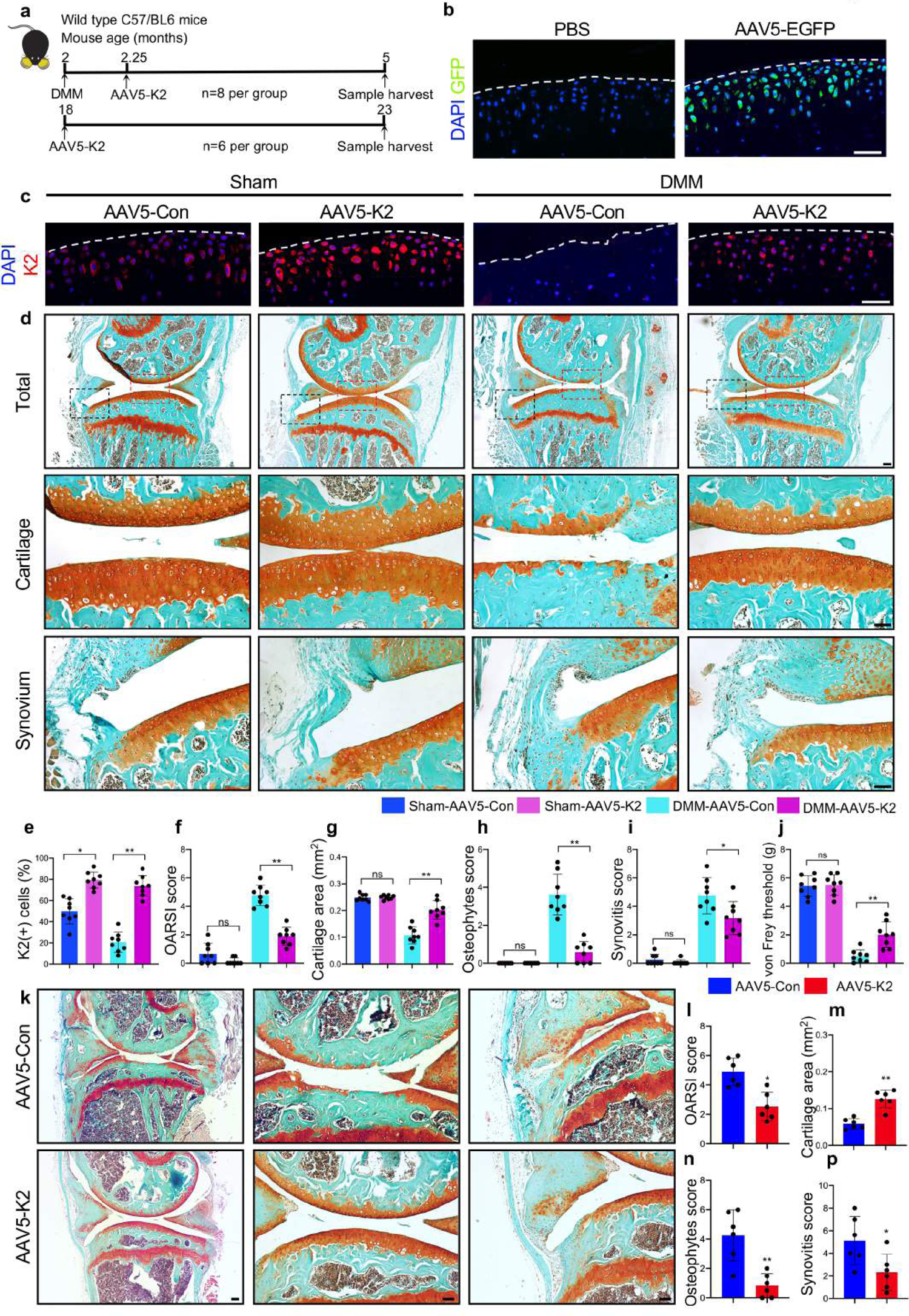
Intraarticular injection of Kindlin-2-expressing adeno-associated virus protects against development of aging- and DMM-induced OA in mice. (**a**) A schematic diagram illustrating the experimental design. (**b**) GFP signals were strongly detected in articular cartilage at 3 weeks after intra-articular injection of AVV5-EGFP. Scale bar, 50 μm. (**c**) IF staining of Kindlin-2 in sham and DMM joint sections from mice intraarticularly injected with AAV5-Con (empty) or AAV5-K2. Scale bar, 50 μm. (**d**) Representative SO&FG staining images of sham and DMM joint sections (upper panels). Black dashed boxes indicate enlarged images of articular cartilage (middle panels). Red dashed boxes indicate synovium (lower panels). Scale bar: 50 μm. (**e**) Percentage of Kindlin-2-positive cells in articular cartilage. *N* = 8 per group. (**f**-**i**) OARSI score (f), cartilage area (g), osteophyte score (h) and synovitis score (i) were analyzed using histological sections. *N* = 8 per group. (j) Quantitative analysis of von Frey threshold (g). (**k**) Representative SO&FG-stained joint sections from aged mice intra-articular injected with AAV5-Con or AAV5-K2. Scale bar, 50 μm. (**l**-**p**) OARSI score (l), cartilage area (m), osteophyte score (n) and synovitis score (o) were analyzed. *N* = 6 per group. All data are the mean ± s.d. \**P* < 0.05, \*\**P* < 0.01. Student’s *t*-test and one-way ANOVA with post hoc test were performed.

## Discussion

In this study, we present the first demonstration that the focal adhesion protein Kindlin-2 maintains the articular chondrocyte anabolism to preserve integrity of the articular cartilage. We demonstrate that Kindlin-2 acts as a critical intrinsic inhibitor of Runx2 expression through inactivation of Stat3 in articular chondrocytes. We show that Kindlin-2 deletion in chondrocytes causes striking and spontaneous OA-like phenotypes, including a progressive loss of the articular cartilage, osteophyte outgrowth, synovial hyperplasia and pain, which highly mimic major features of the aging-associated OA in humans. We show that Kindlin-2 loss accelerates progression of instability-induced OA in adult mice. In both mouse and OA articular cartilage, chondrocytes display reduced expression of Kindlin-2 and increased expression of p-Stat3 and Runx2 proteins. Of translational significance, intraarticular injection of Kindlin-2-expressing AAV5 ameliorates aging-related and DMM-induced OA lesions in mice. Importantly, we provide strong evidence that Kindlin-2 deficiency causes OA by up-regulating Runx2 through promotion of activation and nuclear translocation of Stat3 in articular chondrocytes. These findings highlight a requirement to determine whether abnormalities in expression and/or activity of Kindlin-2, Stat3 and Runx2 in articular chondrocytes play important roles in human OA initiation, development and progression. We may define a useful therapeutic target for OA.

Several studies reported that activation of Stat3 is involved in promotion of OA initiation and progression (*15–17*). However, physiological signals that inhibits Stat3 activation in articular chondrocytes are unclear. In the present study, we identify Kindlin-2 as an intrinsic and potent inhibitor of the Stat3 activation in articular chondrocytes and plays an important role in maintaining the anabolic status of articular cartilage. The loss of Kindlin-2 largely activates Stat3 by increasing its phosphorylation at Y705, while overexpression of Kindlin-2 exerts an opposite effect. siRNA knockdown of Stat3 in chondrocytes reverses the catabolic gene expression pattern caused by Kindlin-2 loss. Interestingly, we find that expression of p-Jak2, a canonical activator of Stat3, is not markedly altered in Kindlin-2 deficient chondrocytes, suggesting that Kindlin-2 loss induced Stat3 activation is not through Jak2 activation. Most importantly, systemic inhibition of Stat3 activation by Stattic or genetic deletion of Stat3 in chondrocytes in mice largely corrects OA lesions, such as cartilage degeneration and osteophyte formation, caused by Kindlin-2 deficiency. These findings, along with our observation that Stat3 is greatly activated in the damaged OA articular cartilage in patients, suggest that Stat3 activation may play an important role in the pathogenesis of human OA. This requires further investigation.

Cumulative evidence points to aberrant accumulation of Runx2 protein in articular chondrocytes being a major player in promoting OA initiation and progression. The loss of Runx2 in chondrocytes provides a significant protection against initiation and progression of instability-induced OA in genetic mouse models, while gain of function of Runx2 stimulates OA development in multiple genetic mouse models (*4, 10-14, 47, 48*). Therefore, it is critical to keep expression of Runx2 in articular chondrocytes under control to preserve integrity of the articular cartilage. Importantly, in the present study, we identify Kindlin-2 as a major player in this respect. We provide multiple lines of molecular and genetic evidence supporting that Kindlin-2 inhibits expression of Runx2 by suppressing Stat3 actions in articular chondrocytes, First, p-Stat3 and Runx2 are in parallel up-regulated in chondrocytes by Kindlin-2 loss in vitro and in cartilage as well as in the aged or damaged articular cartilage of mice and humans with OA. Second, loss of function of Stat3 decreases expression of Runx2 and reverses the chondrocyte catabolic phenotype caused by Kindlin-2 loss in vitro. Third and most importantly, genetic deletion of Stat3 in chondrocytes abolishes Runx2 accumulation and limits OA lesions caused by Kindlin-2 loss in mice, while Runx2 ablation in chondrocytes largely reverses OA lesions caused by Kindlin-2 deletion without down-regulating p-Stat3 in chondrocytes.

Our results of the present study suggest that Kindlin-2 deficiency activates Stat3 by at least in part stimulation of ROS overproduction in chondrocytes. Kindlin-2 deletion results in overproduction of ROS in chondrocyte in vitro and in cartilage. The levels of mitochondrial ROS are also elevated in articular chondrocytes in mouse and human OA cartilage samples. H_2_O_2_ increases the level of p-Stat3 in ATDC5 cells. The ROS scavenger NAC blocks Stat3 activation and up-regulation of Runx2 and ECM-degrading enzymes caused by Kindlin-2 knockdown. Studies from literature also point to a link between excessive mitochondrial ROS accumulation and OA pathogenesis (*49*). While these observations suggest that Kindlin-2 deficiency causes OA partially through up-regulation of ROS in chondrocytes, how Kindlin-2 loss promotes mitochondrial ROS accumulation in chondrocytes remains unclear. It is known that mitochondrion is the primary source for intracellular ROS (*46*). We find that a significant fraction of Kindlin-2 protein exists in mitochondrion of chondrocytes. Our recently published study showed that Kindlin-2 is detected in mitochondrion of lung cancer cells (*34*). A more recent study revealed that suppression of Kindlin-2 mitochondrial translocation and its interaction with pyrroline-5-carboxylate reductase 1 by deletion of Pinch1, another focal adhesion protein, resulted in ROS overaccumulation in lung adenocarcinoma cells (*35*). In addition to Stat3 activation, the loss of Kindlin-2 also promotes Stat3 nuclear translocation in chondrocytes with possible mechanism involving interactions of both factors.

Interestingly, systemic inhibition or genetic deletion of Stat3 in chondrocytes attenuates the cartilage loss and osteophyte outgrowth, but not the synovial hyperplasia, caused by Kindlin-2 deficiency. This suggests that Kindlin-2 deficiency causes synovial hyperplasia not through activation of Stat3. It should be noted that the expression levels of Kindlin-2 protein are comparable in synoviums of control and cKO mice. This is consistent with the fact that *Aggrecan^CreERT2^* is not active in the synovium (Supplementary Figure 2a) (*50*). Thus, the synovial hyperplasia observed in cKO mice is not due to Kindlin-2 deficiency in this tissue.

Loss of β1 integrin activation was reported to disturb chondrocyte cytokinesis, motility and survival, leading to accelerated terminal differentiation of articular chondrocytes (*51*)(*52*). In the present study, we find that loss of Kindlin-2 impairs β1 integrin activation in articular chondrocytes without activating the MAPK pathway. This result is consistent with a previous report showing a normal MAPK activation in β1 integrin-deficient chondrocytes (*52*). In this study, we demonstrate that deletion of Kindlin-2 impairs the attachment and spreading of articular chondrocytes on collagen II coated surfaces. Thus, impaired activation of β1 integrin could partially contribute to the enhanced chondrocyte hypertrophic differentiation and catabolism as well as OA lesions in cKO mice.

It is worthwhile to point out that the spontaneous OA mouse model generated by inducible deletion of Kindlin-2 in adult mice developed in this study is an invaluable tool for OA study in the field. After TM injection, cKO mice develop spontaneous OA phenotypes over time with a 100% penetration (with greater than 100 mice). More importantly, the OA phenotypes highly photocopy those of human OA, including progressive cartilage degeneration, osteophyte formation, synovial hyperplasia and pain. Another advantage of this spontaneous OA mouse model over the DMM OA model is that the former takes a slower process to induce OA initiation, development and progression, which is similar to the pathological process of human OA, a chronic degenerative disease. Furthermore, this model avoids the trouble of the DMM surgery; the latter also creates greater experimental variations.

A large number of elderly people suffer from OA worldwide. However, there are currently no FDA-approved OA treatments or effective interventions to limit OA development and progression. The pathway comprising of Kindlin-2, Stat3 and Runx2 defined in this study may be a useful target for the intervention and treatment of OA. Notably, we provide convincing evidence that intraarticular injection of Kindlin-2-expressing AAV limits progression of aging- and instability-induced OA in mice.

We acknowledge that this study has several limitations. First, while our results show significant limitation of both aging- and instability-induced OA lesions by a single intracellular injection of Kindlin-2 AAV5 in mice, its long-term protective effect against OA remains unclear. This needs to be determined by performing time-course experiments. If the effect of single injection turns out to be unsustainable, we will need to determine whether multiple injections and/or reformulation, for example, by slow release methods, can extend the effectiveness of treatment. Second, in this study, we did not determine whether this injection regimen of AAV-Kindlin-2 will have a similar protective effect against OA development and progression in primates or humans. We plan to perform these experiments on primates in our future study. Third, while expression of Kindlin-2 in articular chondrocytes is down-regulated in mouse and human OA cartilage samples, upstream factors responsible for this down-regulation remain to be determined in future study. Collectively, Kindlin-2 plays a central role in maintaining articular chondrocyte homeostasis and preserving integrity of the articular cartilage and provides a protection against OA.

## Materials and Methods

### Human cartilage samples

Human knee joint cartilage samples were collected from OA patients from the total joint replacement surgeries, following informed written patient consent with approval from The Ethics Committee of Tongji Medical College, Huazhong University of Science and Technology (No. [2020](*43, 44, 53*) IEC-J (565)). Cartilages were excised from tibial plateau and femoral condyles of 8 patients undergoing total knee replacement surgery. Cartilage samples were fixed in 4% paraformaldehyde, decalcified in 15% EDTA and paraffin embedded for further histological and immunofluorescent (IF) analysis.

### Animal studies

Generation of *Kindlin-2^fl/fl^* mice was described (*41*). The *Aggrecan^CreERT2^* mice and floxed Runx2 mice (*Runx2 ^fl/fl^*) were described (*50, 54*). Floxed Stat3 mice (*Stat3^fl/fl^*)(*55*) were kindly provided by Dr. Xin-Yuan Fu of Indiana University School of Medicine. We bred *Kindlin-2^fl/fl^* mice with the *Aggrecan^CreERT2^* knock-in mice to generate the inducible conditional Kindlin-2 knockout mice (*Kindlin-2^fl/fl^; Aggrecan^CreERT2^*). To delete Kindlin-2 expression, mice were administrated with five daily peritoneal injections of tamoxifen (TM, Sigma T5648) at the dosage 100 mg/kg body weight. This TM regimen dramatically reduced Kindlin-2 protein expression in knee joint articular chondrocytes (Figure 2h). *Kindlin-2^fl/fl^; Aggrecan-^Cre/ERT2^* mice treated with corn oil were used as control groups in this study. We crossed the *K2^fl/fl^; Aggrecan^CreERT2^* (cKO) mice with either floxed Stat3 mice (*Stat3^fl/fl^*) or floxed Runx2 mice (*Runx2^fl/fl^*) to generate *K2^fl/fl^; Stat3^fl/fl^; Aggrecan^CreERT2^* mice (^KS^dKO) and *K2^fl/fl^; Runx2^fl/fl^; Aggrecan^CreERT2^* mice (^KR^dKO), respectively. The intraarticular injection of AAV was performed as we previously described (*11*). DMM surgery was performed in the right knees of mice for OA induction according to our previously established protocol (*56*). To minimize use of animals, only male mice were used for experiments in this study. Mice with a C57BL/6 genetic background were used for the experiments in this study. All research protocols in this study were approved by the Institutional Animal Care and Use Committees (IACUC) of Southern University of Science and Technology.

### Micro-computerized tomography

Knee joints were subjected to micro-computerized tomography (μCT) according to our previously established protocol (*11*). Briefly, we used a Skyscan scanner 1276 high-resolution μCT scanner (Bruker, Aartselaar, Belgium) with 60 kVp source and 100 μAmp current for formalin-fixed mouse knee joints with a resolution of 10 μm. The scanned images from each group were evaluated at the same thresholds to allow three-dimensional structural rendering of each sample.

### Animal behavioral tests

Testing for mechanical allodynia (von Frey sensitivity) was performed essentially according to a method previously described (*57*) Before the von Frey test, we allowed animals to adapt to the environment, including an elevated mesh platform for 15 minutes. A calibrated set of von Frey filaments (Stoelting, Wood Dale, IL) was used to poke from below to the hind paw to calculate the 50% force withdrawal threshold using an iterative approach. The tests were performed in a blind manner that the investigator is not aware of the identification of animals as well as the study groups.

### Isolation of primary articular chondrocytes

The isolation and culture of primary chondrocytes from adult articular cartilage were modified from a previously described protocol (*58*). Two-month-old male mice were sacrificed and the hindlimbs were dissected. The hindlimbs were washed three times with PBS and then skin and soft tissues were removed from bones using scissors and pincer in a sterile flow hood. Articular cartilages on femoral condyles and tibial plateau were peeled off using a blunt-ended forceps and surrounding synovial layer and tendons were carefully removed under a stereo microscope. Then, the isolated cartilage was crushed into small pieces and digested in 0.25% trypsin-EDTA solution for 20 mins, followed by an overnight digestion in collagenase II solution (1mg/ml). The released cells from cartilage pieces were washed by PBS and cultured in 60 mm plates with Dulbecco‘s Modified Eagle Medium/F12 medium supplemented with 10% fetal bovine serum, 1% glutamine and 1% penicillin and streptomycin at 37℃ with 5% CO_2_ for further in vitro experiments. To induce deletion of Kindlin-2 in primary articular chondrocytes of *Kindlin-2^fl/fl^; Aggrecan^CreERT2^* mice in vitro, 4-Hydroxytamoxifen Ready Made Solution (Sigma, SML1666) was added into the culture medium (1:1000) for 48 hours. Vehicle (Ethanol: isopropanol (95:5)) was used as control. The expression levels of Kindlin-2, Stat3, p-Stat3 and Runx2 were analyzed by western blotting and IF staining.

### siRNA knockdown experiments

ATDC5 cells were transfected with the indicated siRNAs. 48h after transfection, protein extracts were isolated from cells and subjected to western blot analyses with indicated antibodies. The sequences of siRNA primers used in this study are summarized in Supplementary Table 1.

### Isolation of mitochondrial and cytosolic fractions

Mitochondrial and cytosolic fractions were isolated using a Mitochondria Isolation Kit (Thermo Fisher Scientific, Cat #898874) according to our previously established protocol (*34*). Briefly, cultured primary articular chondrocytes were harvested by centrifuge at 850g for 2 minutes and resuspended in a 2.0-ml microcentrifuge tube. Then, Reagent A, B and C was sequentially added to the cells according to manufacturer’s instructions. The cells were centrifuged at 700g for 10 minutes at 4°C to remove the nuclei. The supernatants were further centrifuged at 12,000g for 15 minutes at 4°C and then the supernatants (cytosolic fractions) and pellets (mitochondrial fractions) were collected.

### Measurement of ROS

The measurement of ROS in homogenized articular cartilage was performed using a 6-chloromethyl-2‘,7’-dichlorodihydrofluorescein diacetate, acetyl ester; (CM-H_2_DCFDA)-based commercial kit (GENMED, GMS10016.3). Briefly, articular cartilages were isolated from the hindlimbs of mice and homogenized using BioPulverizer system (Cat# 59012N). The protein concentrations of homogenized tissues were determined using Pierce™ BCA Protein Assay Kit (Thermo Fisher, Cat# 23225). The homogenized tissues were then incubated with CM-H_2_DCFDA probe at 37 °C for 20 min with protection from light. The fluorescence intensity was measured and analyzed by EnSpire system (PerkinElmer). The relative fluorescence intensity was taken as the average values from three repeated experiments. The mitochondrial ROS levels were measured using the MitoSOX red Mitochondrial Superoxide Indicator (Thermo Fisher, Cat# M36008) following the manufacturer’s instruction. The MitoSOX reagents were prepared at a 5 μM working concentration in HBSS/Ca/Mg buffer. For the detection of mitochondrial ROS in cultured cells, cells were washed 3 times by PBS and then loaded with 5 μM MitoSOX reagent and incubated at 37 °C for 10 min under protection of light. For the measurement of ROS levels in articular cartilage, paraformaldehyde-fixed, sections (5 μm) were incubated with 5 μM MitoSOX Red for 15 min at room temperature under protection of light as previously described (*59, 60*). Nuclei were stained with DAPI. the fluorescence signals were analyzed by SP8 Leica confocal microscopy (excitation wavelength 510 nm, emission wavelength 580 nm). Human cartilage samples were fixed in Methacarn fixative solution (10% glacial acetic acid, 30% trichloromethane and 60% methanol) at 4 °C overnight and decalcified in 15% EDTA (without PFA) in PBS. OxyIHC staining was performed according the Millipore OxyIHCTM oxidative stress detection kit protocol (Millipore, S7450).

### RNA sequencing analysis

Total RNA was extracted from homogenized articular cartilage tissues of control and cKO mice at 5 months after TM injections using a TransZol Up Plus RNA Kit (ER501-01; Transgen, China) and 3 μg RNA per sample was used as input material for the RNA sample preparations. Sequencing libraries were generated using NEBNext R UltraTM RNA Library Prep Kit (Illunina, NEB, United States) and the library quality was assessed on the Agilent Bioanalyzer 2100 system. After cluster generation, the library preparations were sequenced on an Illumina Hiseq platform and 150 bp paired-end reads were generated. After quality control, reads mapping to the reference genome and quantification of gene expression level, KEGG enrichment analyses were performed by using the cluster Profiler R package.

### Western blot analyses

Western blot analysis was performed as previously described (*61*). Briefly, protein extracts were fractionated on a 10% SDS-PAGE gel and transferred onto nitrocellulose membranes (Schleicher & Schuell, Keene, NH). The membrane was blocked in 5% nonfat milk in Tris-buffered saline/Tween 20 buffer and probed with primary antibodies, followed by incubation with secondary antibodies conjugated with horseradish peroxidase, and then visualized using a Western Blotting Detection Kit (GE Healthcare, cat#: RPN2106). Antibodies used in this study are listed in Supplementary Table 2.

### Histology, immunofluorescence and confocal analysis

Knee joint tissues were fixed in 4% paraformaldehyde, decalcified, dehydrated, and embedded in paraffin. Serial sections (5-μm thick) were cut and stained with Safranin O & Fast Green (Solarbio, Cat#G1371)/hematoxylin and eosin (Thermo Fisher Cat#7211&7111) for morphological analysis according to manufacturer’s instructions. For IF staining, 5-μm sections were permeabilized with 0.2% Triton X-100, blocked with 2% bovine serum albumin (BSA) for 1h and then incubated with primary antibodies (Supplemental Table 2) overnight at 4°C. After washing, the sections were incubated with anti-rabbit Alexa Fluor 488 (Invitrogen) or anti-mouse Alexa Fluor 568 (Invitrogen) secondary antibodies (1:400) for 1h at room temperature. The fluorescent signals in articular cartilage areas were determined using a confocal microscope (Leica SP8 Confocal Microsystems).

### Quantitative real-time PCR analysis

Total RNA was extracted from cells with TriPure Isolation Reagent (Sigma-Aldrich) as previously described (*62*). Reverse transcription to cDNA was prepared by using Superscript II reverse transcriptase (Invitrogen) and oligo(dT) primers. Real-time PCR was performed using SYBR ® Premix Ex Taq™ II with an ABI 7500 QPCR System. Mouse *Gapdh* mRNA levels were used as an internal control of the target mRNAs. Normalization and fold changes were calculated using the ΔΔCt method. Primer sets are listed in Supplementary Table 3.

## Statistical Analysis

The sample size for each experiment was determined based on our previous experience. Animals used in experiments of this study were randomly grouped. IF and histology were performed and analyzed in a double-blinded way. Statistical analyses were completed using the Prism GraphPad. The two-tailed unpaired Student’s *t* test (two groups) and one-way ANOVA (multiple groups), followed by Tukey’s post-hoc test, were used. Results are expressed as mean ± standard deviation (s.d.), as indicated in the Figure Legends. Differences with *P* < 0.05 were considered statistically significant.

## Data availability

All data generated for this study are available from the corresponding authors upon reasonable request.

## Acknowledgments

The authors acknowledge the assistance of Core Research Facilities of Southern University of Science and Technology. This work was supported, in part, by the National Key Research and Development Program of China Grants (2019YFA0906004), the National Natural Science Foundation of China Grants (81991513, 82022047, 81630066, 81870532, 82972100), the Guangdong Provincial Science and Technology Innovation Council Grant (2017B030301018), and the Science and Technology Innovation Commission of Shenzhen Municipal Government Grants (JCYJ20180302174246105, JCYJ20180302174117738 and KQJSCX20180319114434843).

## Author contributions

Study design: GX, XW, YL and HC. Study conduct and data collection: XW, YL, SC, CZ, XF, CT, JL, JH, WT, HT, XB and GX. Data analysis: XW, YL and GX. Data interpretation: GX, XW, CL, ZS, XB and DC. Drafting the manuscript: GX and XW. XW, HC and GX take the responsibility for the integrity of the data analysis.

