## Supplementary information for "Kindlin-2 preserves integrity of the articular cartilage to protect against osteoarthritis"

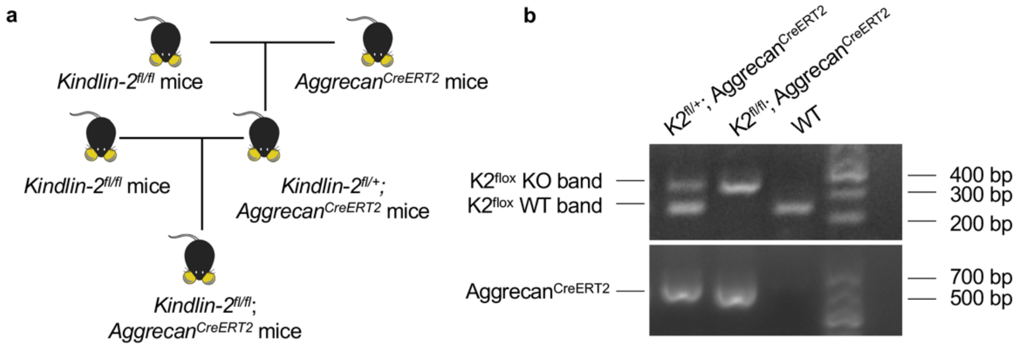


**Supplementary Figure 1. Breeding strategy and PCR genotyping.** (**a**) Breeding strategy. (**b**) PCR genotyping using tail DNA. K2$^{flox}$ KO, ~300bp; K2$^{flox}$ WT, ~200bp; Aggrecan$^{CreERT2}$, ~650bp. Primer sets are listed in Supplementary Table 4.

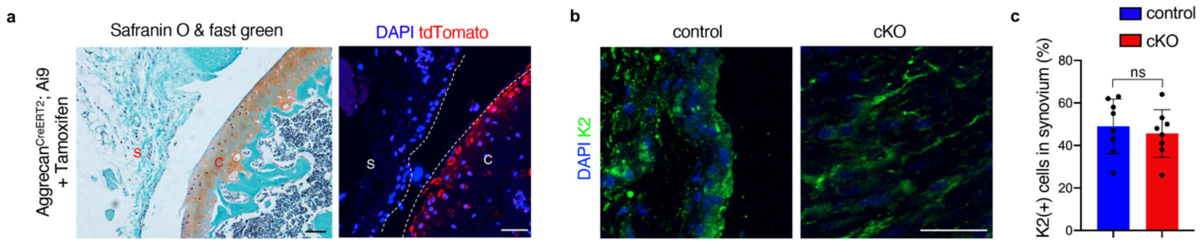


**Supplementary Figure 2.** ***Aggrecan$^{CreERT2}$*** **activity and Kindlin-2 expression in synovium.** (**a**) Representative images of safranin O & fast green and fluorescent images showing the tdTomato expression in knee joint sections *from Aggrecan$^{CreERT2}$; Ai9* mice at 4 weeks after tamoxifen injections. S: synovium; C: cartilage. Scale bar: 50 μm. (**b**) Immunofluorescent (IF) staining for expression of Kindlin-2 in control and cKO synovium. Scale bar: 50 μm. (**c**) Percentage of Kindlin-2-positive cells in control and cKO synovium. Results are expressed as mean ± standard deviation (s.d.). ns: not significant. $N = 8$ mice per group.

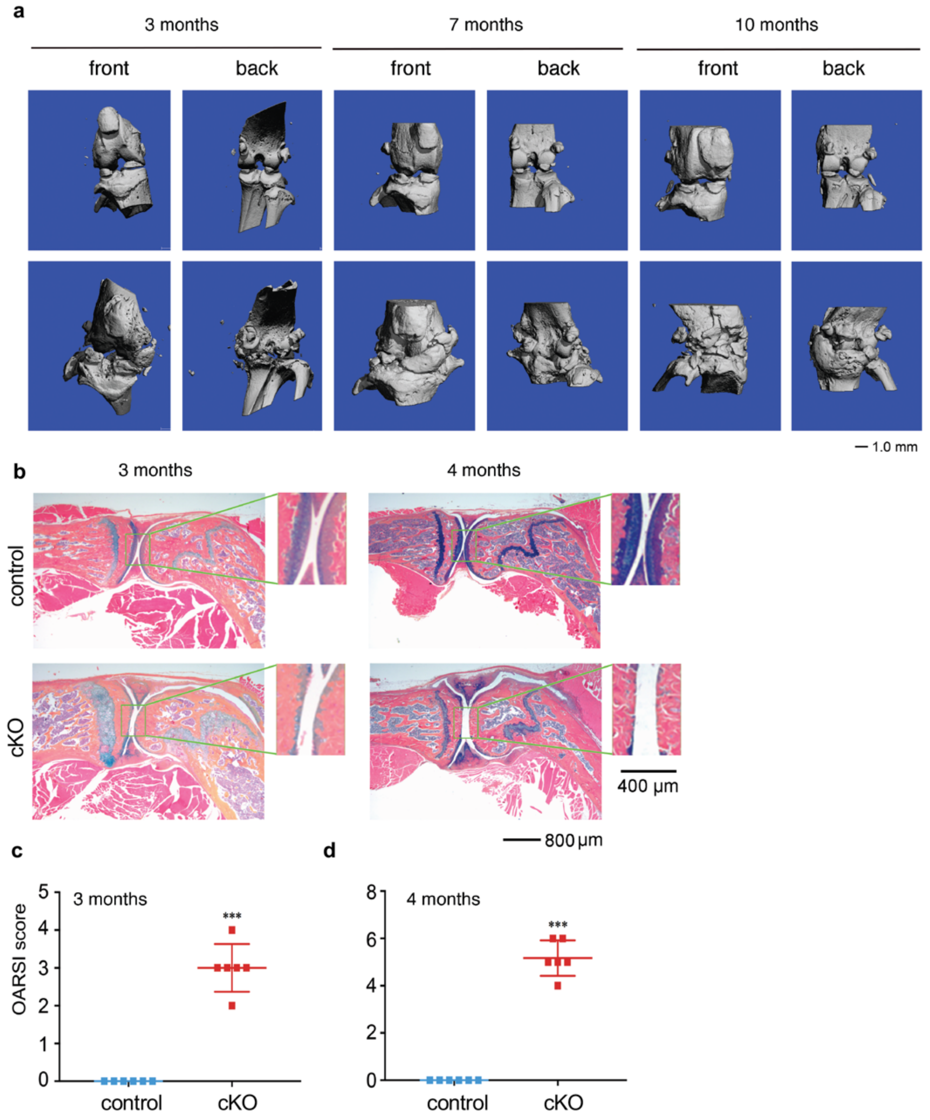


**Supplementary Figure 3. Knee joint OA progression in cKO mice.** (**a**) Three-dimensional reconstruction form μCT scans of knee joints. At 3, 7 and 10 months after TM injection, knee joints of control and cKO male mice were subjected to μCT analysis. Scale bar, 1.0 mm. (**b**) Alcian blue/hematoxylin and eosin stain. At 3 and 4 months after TM injection, knee joint sections of control and cKO male mice were subjected to alcian blue/hematoxylin and eosin stain. Representative images from the two genotypes at the indicated ages are shown. Representative lower magnification images (left panels) and higher magnification images of the boxed areas (right panels) are shown. Scale bar, 800 μm and 400 μm as indicated. (**c,d**) OARSI score. $N$ = 6 mice per group for both control and cKO mice. ***$P < 0.001$, versus control, Student's $t$ test. Results are expressed as mean ± standard deviation (s.d.).

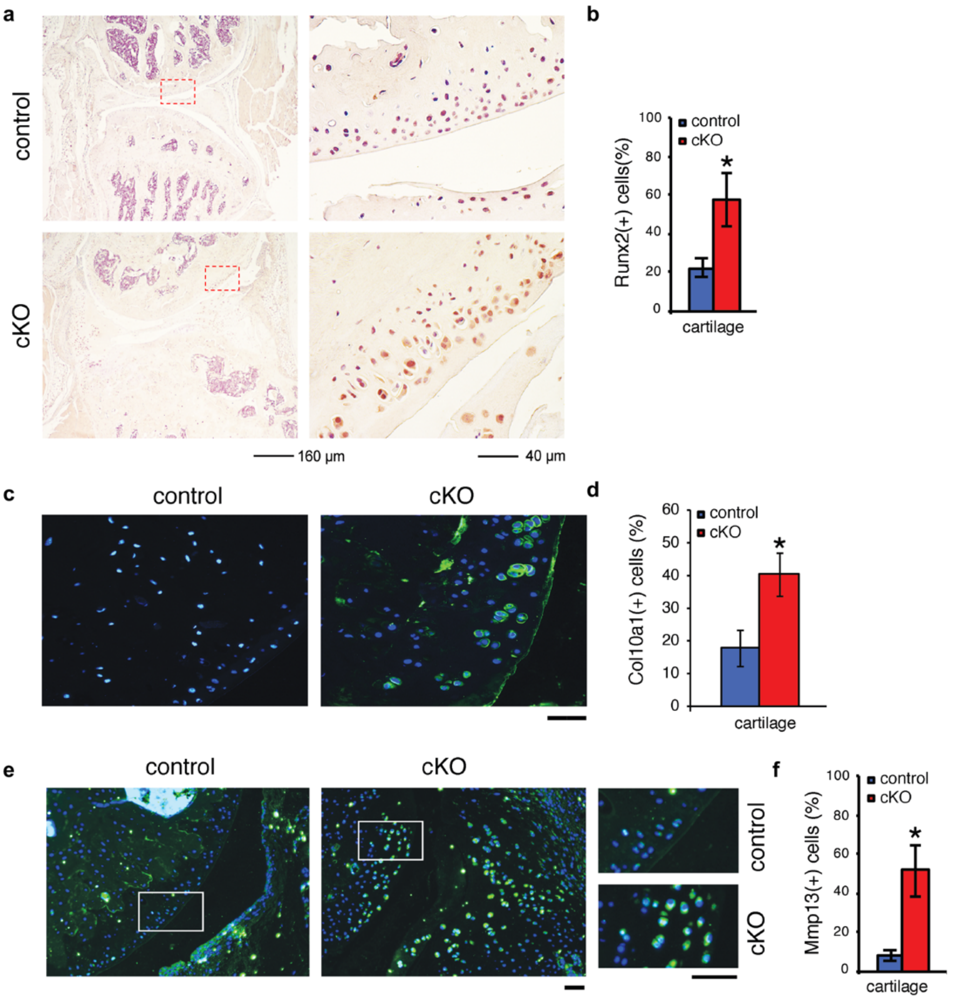


**Supplementary Figure 4. Kindlin-2 loss promotes chondrocyte hypertrophic and catabolic phenotypes in articular cartilage.** (**a,b**) Immunohistochemical (IHC) staining. Three months after TM injection, knee joint sections of control and cKO mice were subjected to IHC staining with an antibody against Runx2. Representative images from the two genotypes showing increased expression Runx2 in cKO articular cartilage. Scale bar, 160 μm or 40 μm as indicated. Quantitative data (b). (**c-f**) IF staining. Three months after TM injection, knee joint sections from control and cKO mice were stained with antibodies against Col10a1 (c) and Mmp13 (e). Results show dramatic chondrocyte hypertrophy together with increased expression of Col10a1 and Mmp13. Scale bar, 20 μm. Quantitative data (d,f). $N = 9$-10 mice for each group. $^*P < 0.05$, versus control, Student's *t* test. Results are expressed as mean ± standard deviation (s.d.).

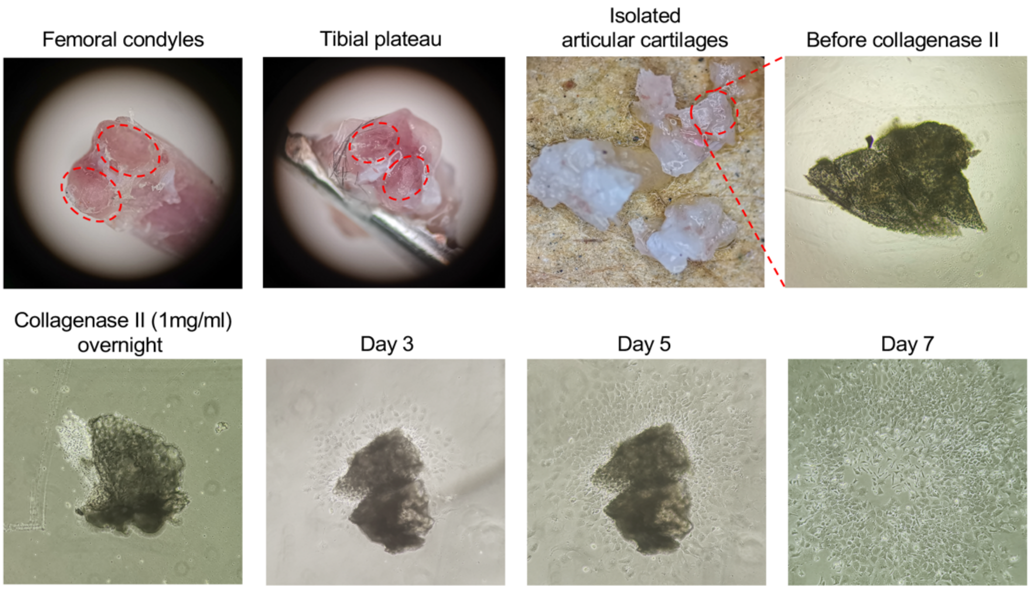


**Supplementary Figure 5. Isolation of primary chondrocytes from adult *K2$^{fl/fl}$; Aggrecan$^{CreERT2}$* male mice.** Two-month-old *K2$^{fl/fl}$; Aggrecan$^{CreERT2}$* male mice were sacrificed and the hindlimbs were dissected. Articular cartilages on femoral condyles and tibial plateau were peeled off using a blunt-ended forceps and surrounding synovial layer and tendons were carefully removed under a stereo microscope. The isolated cartilage was then crushed into small pieces and digested in 0.25% trypsin-EDTA solution for 20 mins, followed by an overnight digestion in collagenase II solution (1mg/ml). The released cells from digestion were washed by PBS and cultured in 60 mm plates with Dulbecco's Modified Eagle Medium/F12 medium supplemented with 10% fetal bovine serum, 1% glutamine and 1% penicillin and streptomycin at 37°C with 5% $CO_2$ for further in vitro experiments.

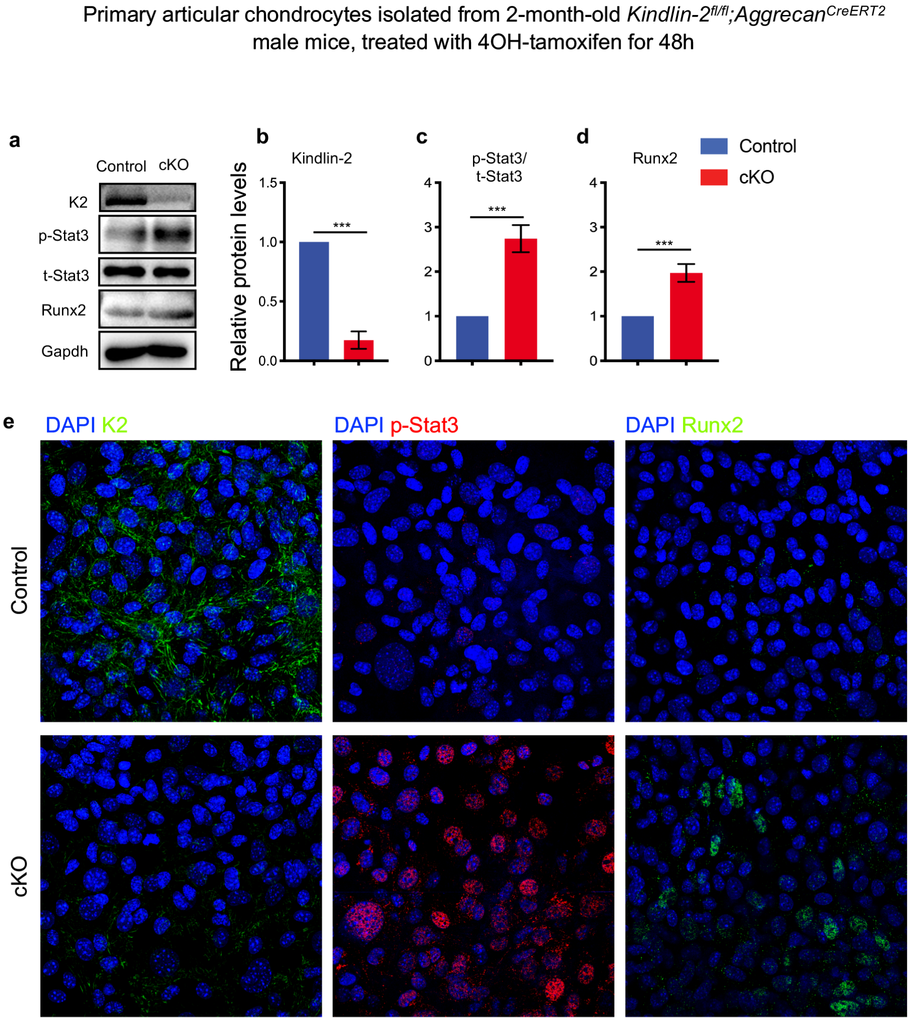


**Supplementary Figure 6. Kindlin-2 deletion increases the levels of p-Stat3 and Runx2 proteins in primary chondrocytes.** (**a**) Primary articular chondrocytes were isolated from 2-month-old *K2*$^{fl/fl}$*; Aggrecan*$^{CreERT2}$ male mice and cultured in 6-well plates. The cells were then treated with 4-hydroxytamoxifen (cKO) and vehicle (control) for 48h. The expression levels of Kindlin-2, t-Stat3, p-Stat3 and Runx2 were analyzed by western blotting and IF staining. (**b-d**) Quantitative data of western blotting analysis of Kindlin-2 (b), p-Stat3/t–Stat3 (c) and Runx2 (d). Results are expressed as mean ± standard deviation (s.d.). Experiments were repeated three times independently. ***$P$ < 0.001. (**e**) IF staining of Kindlin-2, p-Stat3 and Ruxn2. Nuclei were labelled by DAPI.

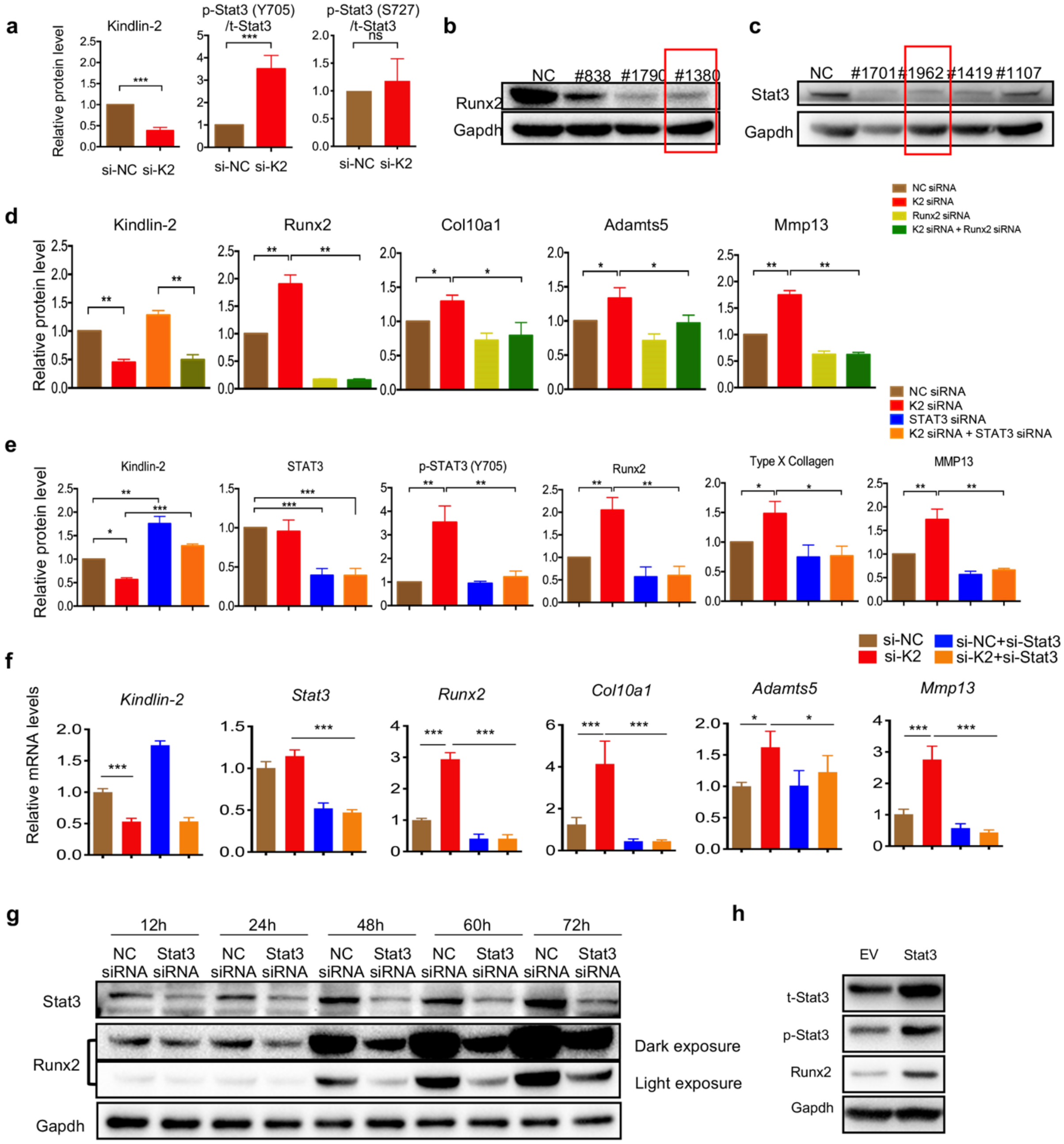

**Supplementary Figure 7. siRNA knockdown / overexpression of Stat3 and Runx2 in chondrocytes.** (**a**) Quantitative data for Figure 4e. (**b, c**) siRNA knockdown of Runx2 and Stat3 in ATDC5 cells. siRNA #1380 (for Runx2) and #1962 (for Stat3) were used for subsequent experiments. (**d**) Quantitative data for Figure 4u. (**e**) Quantitative data for Figure 4v. (**f**) ATDC5 cells were transfected with si-NC or si-K2 with and without si-Stat3, followed by qPCR analysis. (**g**) ATDC5 cells were transfected with si-NC or si-Stat3, followed by western blotting for expression of Stat3 and Runx2. (**h**) ATDC5 cells were transfected with empty vector (EV) and Stat3 expression vector (Stat3), followed by western blotting for expression of t-Stat3, p-Stat3 and Runx2. Results are expressed as mean ± standard deviation (s.d.). $*P < 0.05$; $**P < 0.01$; $***P < 0.001$; ns: not significant.

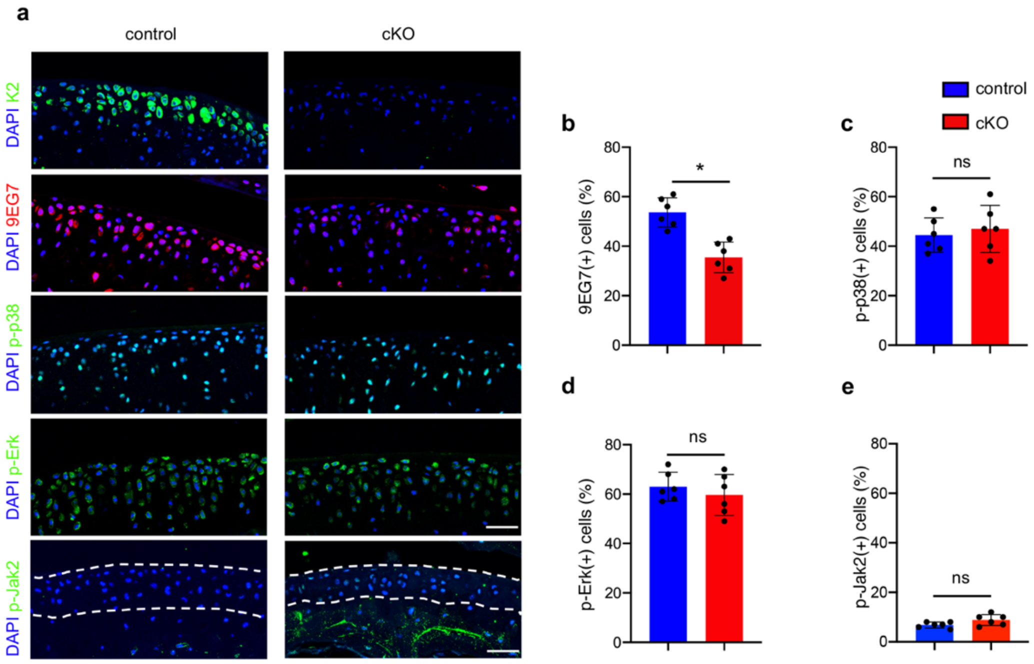


**Supplementary Figure 8. Immunofluorescent analysis of articular chondrocytes in control and Kindlin-2 cKO mice.** (**a**) IF staining for expression of Kindlin-2, 9EG7, p-p38, p-Erk and p-Jak2 in articular chondrocytes from control and cKO mice at 6 months after tamoxifen injections. Scale bar: 50 μm. (b-d) Percentage of positive-stained cells in control and cKO articular chondrocytes. Results are expressed as mean ± standard deviation (s.d.). ns: not significant. *$P < 0.05$. $N = 6$ mice per group.

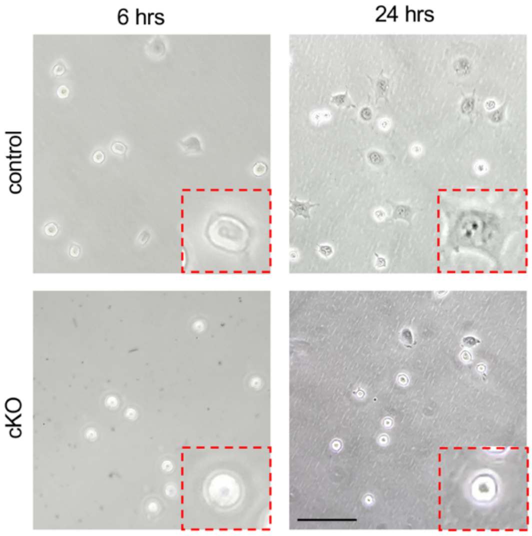


**Supplementary Figure 9. Effects of Kindlin-2 loss on the attachment and spreading of primary articular chondrocytes.** (**a**) Representative images of control and cKO articular chondrocytes attachment and spreading on type II collagen-coated surfaces at 6 and 24 h after seeding. Higher magnification images are shown in red dashed boxes. Scale bar: 50 μm.

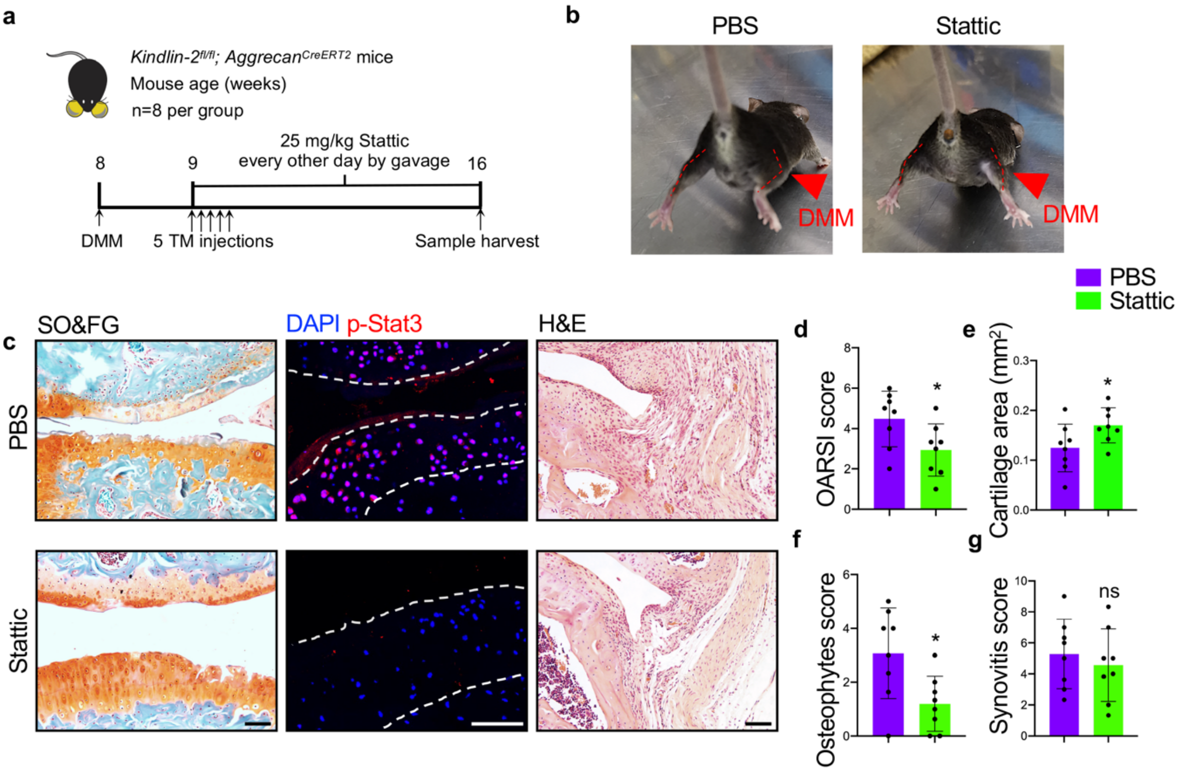


**Supplementary Figure 10. Systemic inhibition of Stat3 activation limits cartilage degeneration and osteophyte formation caused by Kindlin-2 loss.** (**a**) A schematic diagram illustrating the experimental design. (**b**) Stattic improves restriction of movement of knee joint caused by Kindlin-2 loss in DMM mice. (**c**) SO&FG (left panels), IF (middle panels) and H&E (right panels) staining of knee joint sections from cKO mice treated with PBS or Stattic for 2 months after DMM surgery. Scale bar, 50 μm. (**d-g**) OARSI score (d), cartilage area (e), osteophyte score (f) and synovitis score (g) were analyzed using histological sections. $N = 6$. Results are expressed as mean ± standard deviation (s.d.). $^*P < 0.05$.

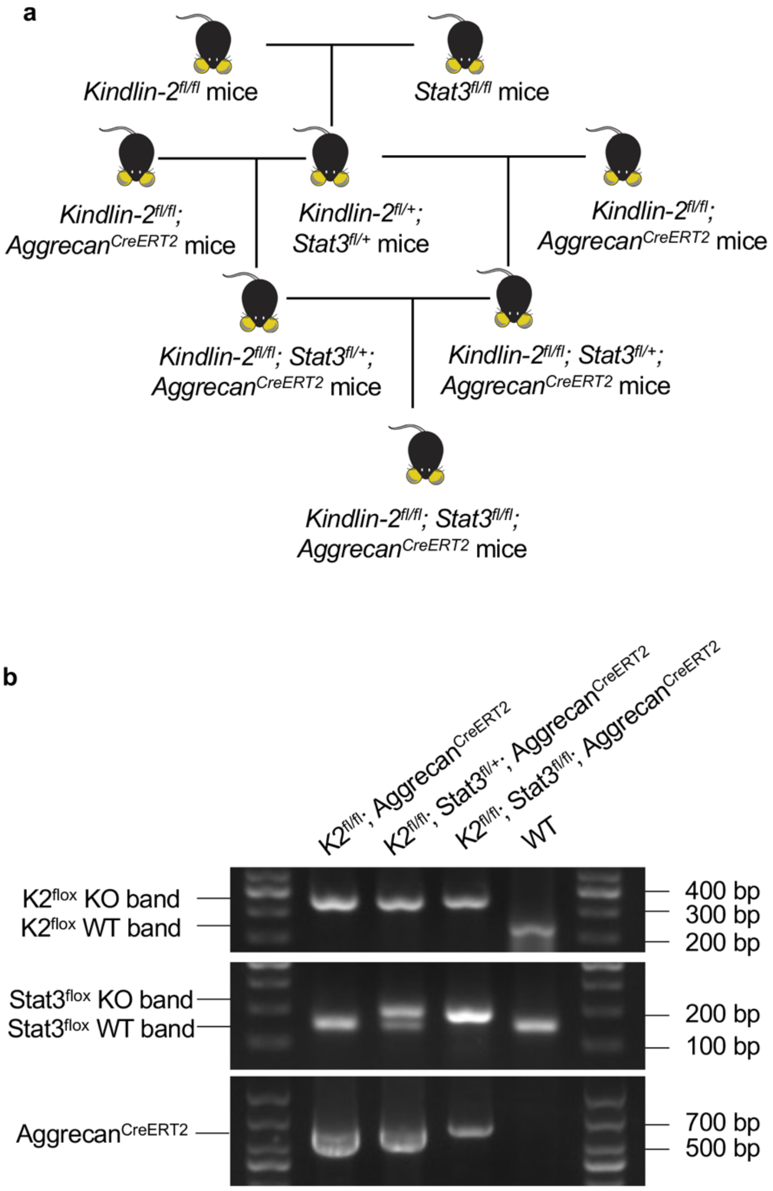


**Supplementary Figure 11. Breeding strategy and PCR genotyping.** (**a**) Breeding strategy for generating the *K2*$^{fl/fl}$*; Stat3*$^{fl/fl}$*; Aggrecan*$^{CreERT2}$ ($^{KS}$dKO) mice. (b) PCR genotyping using tail DNA. K2$^{flox}$ KO, ~300bp; K2$^{flox}$ WT, ~200bp; Stat3$^{flox}$ KO, 187bp; Stat3$^{flox}$ WT, 146bp; Aggrecan$^{CreERT2}$, ~650bp. Primer sets are listed in Supplementary Table 4.

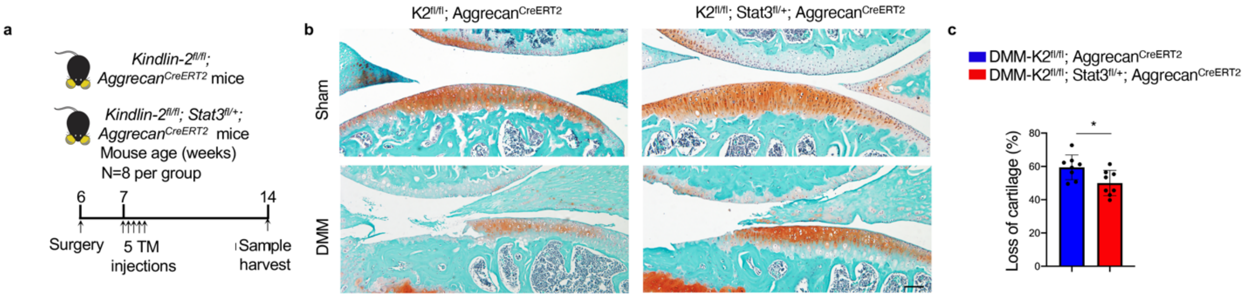


**Supplementary Figure 12. Effect of Stat3 haploinsufficiency in chondrocytes on cartilage loss caused by Kindlin-2 loss.** (**a**) A schematic diagram illustrating the experimental design. (**b**) Safranin O & fast green-stained sections knee joints of *K2$^{fl/fl}$; Aggrecan$^{CreERT2}$* mice and *K2$^{fl/fl}$; Stat3$^{fl/+}$; Aggrecan$^{CreERT2}$* mice subjected to sham or DMM surgeries and tamoxifen injections as described in (a). Scale bar: 50 μm. (**c**) Quantification of cartilage loss. Results are expressed as mean ± standard deviation (s.d.). **P* < 0.05. *N* = 8 mice per group.

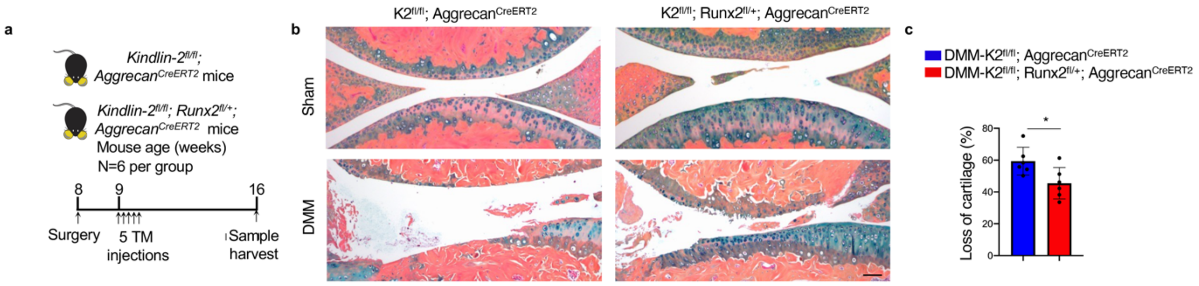


**Supplementary Figure 13. Effect of Runx2 haploinsufficiency in chondrocytes on cartilage loss caused by Kindlin-2 loss.** (**a**) A schematic diagram illustrating the experimental design. (**b**) Alcian blue & orange G-stained sections of knee joints of *K2$^{fl/fl}$; Aggrecan$^{CreERT2}$* mice and *K2$^{fl/fl}$; Runx2$^{fl/+}$; Aggrecan$^{CreERT2}$* mice subjected to sham or DMM surgeries and tamoxifen injections as described in (a). Scale bar: 50 μm. (**c**) Quantification of cartilage loss. Results are expressed as mean ± standard deviation (s.d.). **P* < 0.05. *N* = 6 mice per group.

**Supplementary Table 1: siRNA sequences**

| Name | 5' primer | 3' primer |
|---|---|---|
| Kindlin-2 | GUGGCUAGAUUCCUCAAGATT | UCUUGAGGAAUCUAGCCACTT |
| Stat3 | GGGUCUGGCUAGACAAUAUTT | AUAUUGUCUAGCCAGACCCTT |
| Runx2 | GCAGAAUGGAUGAGUCUGUTT | ACAGACUCAUCCAUUCUGCTT |

**Supplementary Table 2: Antibody information**

| Antibody | Company | Catalog # | Application/Dilution |
|---|---|---|---|
| Kindlin-1 | Sigma | SAB4200465 | IF (1:200) |
| Kindlin-2 | Proteintech | 11453-1-AP | WB (1:1000); IF (1:200) |
| Kindlin-2 | Millipore | MAB2617 | IF (1:200) |
| Kindlin-3 | CST | 13843 | IF (1:200) |
| Runx2 | CST | 12556 | WB (1:1000) |
| Runx2 | Abcam | ab23981 | IHC (1:1000); IF (1:200) |
| Col10a1 | Abcam | ab58632 | WB (1:1000); IHC (1:1000); IF (1:200) |
| Mmp13 | Abcam | ab39012 | WB (1:3000); IHC (1:1000); IF (1:200) |
| Tubulin | CWBIO | CW0098 | WB (1:1000) |
| Vdac | CST | 4661 | WB (1:1000) |
| Stat3 | Abclonal | A19566 | WB (1:1000); IF (1:200) |
| p-Stat3 (Y705) | Abclonal | AP0070 | WB (1:1000); IF (1:200) |
| p-Stat3 (S727) | Abclonal | AP0715 | WB (1:1000) |
| Adamts5 | Abcam | ab41037 | WB (1:500) |
| Gapdh | ZhongShanJinQiao | TA-08 | WB (1:2000) |
| 9EG7 | BD Pharmingen | 553715 | IF (1:200) |
| p-p38 | CST | 4511 | IF (1:200) |
| p-Erk | CST | 9101S | IF (1:200) |
| p-Jak2 | Abclonal | AP0531 | IF (1:200) |

**Supplementary Table 3: Primer information**

| Gene | Forward | Reverse |
|---|---|---|
| *Kindlin-2* | TTCATTCAAGCCTGGCAGTCTCTG | GGCGTCCATCCGAATCAACCTG |
| *Stat3* | TGTCAGATCACATGGGCTAAAT | GGTCGATGATATTGTCTAGCCA |
| *Runx2* | CCTTCAAGGTTGTAGCCCTC | GGAGTAGTTCTCATCATTCCCG |
| *Col10a1* | GAATTTCTGTGCCAGGAAAACC | TTTTCACCTCTTCTTCCCACTC |
| *Adamts5* | GGCAAATGTGTGGACAAAACTA | GAGGTGCAGGGTTATTACAATG |
| *Mmp13* | CTTCCTGATGATGACGTTCAAG | GTCACACTTCTCTGGTGTTTTG |

**Supplementary Table 4: Genotyping primer information**

| Gene | Forward | Reverse |
|---|---|---|
| *Kindlin-2* | 5'-TGTGTTTCAAAGGTACTGGTCA-3' | 5'-ACAATGGTGCTTTGCCTACA-3' |
| *Stat3* | 5'-TTG ACC TGT GCT CCT ACA AAA A-3' | 5'-CCC TAG ATT AGG CCA GCA CA-3' |

| | | |
|---|---|---|
| *Runx2* | 5’-TAA ATC CAG ATG CCC CTG AG-3’ | 5’-TTG AAA CCA TCC ACA GGT GA-3’ |
| Aggrecan CreERT2 | 5’-GAT CTC CGG TAT TGA AAC TCC AGC-3' | 5’-GCT AAA CAT GCT TCA TCG TCGG-3' |
